# Recombination on “non-recombining” sex chromosomes: empirical insights from the blackspotted stickleback

**DOI:** 10.64898/2026.07.23.740300

**Authors:** Zuyao Liu, Karolina Wąchała, Stephan Peischl, Changde Cheng, Mark Kirkpatrick, Catherine L. Peichel, Daniel L. Jeffries

## Abstract

One hallmark of sex chromosome evolution is recombination suppression driven by inversions. The resulting non-recombining sex-linked regions accumulate deleterious mutations and functionally degenerate. The question therefore arises: why does selection not favor the reestablishment of recombination? Two (non-exclusive) explanations for this have been proposed: 1) selection to maintain linkage between the Primary Sex Determination Locus (PSDL) and sexually antagonistic loci - loci with alleles that are beneficial to one sex but detrimental to the other; and 2) the rarity of reversions - structural mutations that reverse the inversion that initially suppressed recombination. Empirical perspectives are sorely needed to evaluate the relevance of these models. Here, we characterised the intermediate stages of sex chromosome evolution in the blackspotted stickleback (*Gasterosteus wheatlandi*). We generated high-quality, phased assemblies and annotations for the X and Y chromosomes, as well as population-level sequencing. Using these data, we inferred distinct recombination loss events and their associated inversions. Most significantly, we find support from multiple lines of evidence for a ∼2 Mb recombination event that likely occurred via double crossover within an existing non-recombining inversion. This result highlights that rare recombination can occur within sex-linked inversions. Simulations suggest that such histories of gene flux may allow for a novel empirical framework, adapted from work on autosomal inversions, to infer evolutionary processes at work within sex-linked non-recombining regions and distinguish between models to explain the maintenance of recombination suppression on sex chromosomes.

**Significance Statement:** Sex chromosomes have evolved countless times across eukaryotes and often lose the ability to recombine, causing them to degenerate over time. Why then, does recombination not restart to purge harmful mutations? By generating a high-quality genome assembly of the blackspotted stickleback, we discovered the strong empirical evidence of a sex chromosome undergoing a double-crossover event within a “non-recombining” region, which reset the degenerative process. This finding challenges the view that sex chromosome decay is a one-way street. Furthermore, using forward evolutionary simulations, we demonstrate that the genetic signatures left by such events could help identify the selective forces acting within non-recombining sex chromosomes, thereby explaining the maintenance of recombination suppression.

## Introduction

Recombination loss is a common feature of sex chromosomes (1) and can link large regions of the genome to the primary sex determination gene. Such regions evolve in unique ways, often becoming hotspots for loci involved in sexual dimorphism and sexual selection (2), disease (3), adaptation (4), and speciation (5–8). The forces that drive the initial suppression of recombination on sex chromosomes have received much attention, and various models have been proposed to explain this phenomenon (see reviews 9, 10). In contrast, much less attention has been given to the long-term maintenance of recombination suppression. Recombination has been lost hundreds of millions of years ago on the sex chromosomes of many taxa, including mammals (∼180 My) (11), birds (∼100 My) (12, 13), and lepidoptera (∼180 My) (14). In these cases, deleterious mutations have accumulated in non-recombining regions due to a reduction in the ability of selection to efficiently remove them (1, 15). This raises the question of why the restoration of recombination has not been favoured by selection in these regions, given that it would allow for the purging of accumulated deleterious load?

Two main mechanisms have been suggested to resist the reestablishment of recombination on sex chromosomes. The first is that non-recombining haplotypes that link the Primary Sex Determination Locus (PDSL) to sexually antagonistic loci (loci with alleles that are beneficial to one sex but detrimental to the other) are fitter than recombinant haplotypes. Thus, recombinants would be maladaptive and unable to fix (16). For example, a recent theory suggests that divergence of gene regulation between the X and Y after recombination loss could result in more male-beneficial, female-detrimental regulation of genes on the Y (or *vice versa* on the W) (17). However, this, and all other theories that invoke adaptive linkage face the same problem: the erosion of Y (or W) chromosome fitness due to continued accumulation of deleterious mutations over time. Theoretical work predicts that this will eventually negate the benefits from sex-linked loci (18–20).

The second mechanism proposed to resist reestablishment of recombination on sex chromosomes is constraint on the required mutations. Recombination loss on sex chromosomes is often driven by inversions (21–24). Theoretically, a second inversion, involving the same breakpoints (a reversion) could restore recombination (17). However, it has been argued that such events are extremely unlikely (25). Debate is ongoing concerning the plausibility of these mechanisms (16), which are not mutually exclusive. What is lacking from this debate is empirical data with which to determine their relative likelihoods.

Stickleback fish in the Gasterosteidae family are a promising system to test such models. There have been several sex chromosome turnovers among sticklebacks (26, 27), and sex chromosomes vary in the timing of recombination suppression (21, 28, 29), making them ideal for studying the early and intermediate stages of sex chromosome evolution. Here, we focus on the blackspotted stickleback, *Gasterosteus wheatlandi*, which is closely related to the well-studied threespine stickleback *G. aculeatus*. These two species share an ancestral XY sex chromosome system on chromosome (Chr) 19 and the oldest nonrecombining region (or “stratum”) with an estimated maximum age of 22 Mya (21, 30). *G. wheatlandi* and *G. aculeatus* diverged approximately 14 Mya (31). Since then, the *G. wheatlandi* Y chromosome has lost recombination along the rest of Chr19 and fused to Chr12. The Chr12 neo-Y chromosome has also since lost recombination along much of its length (27, 30).

Previous work characterising the evolutionary history of the *G. wheatlandi* sex chromosomes aligned short-read sequences to the X chromosome of the *G. aculeatus* genome assembly (30). However, the lack of reference assemblies for the sex chromosomes in *G. wheatlandi* limited the inferences possible concerning the location, timing and mechanisms of recombination suppression, and the content of evolutionary strata. Here we produce a high-quality assembly and annotation for *G. wheatlandi*, including phased assemblies and annotations of its X and Y chromosomes. We use several sequencing and population genetics metrics to characterise divergence and degeneration along the sex chromosome and infer evolutionary strata. In doing so, we discovered strong evidence that at least one recombination event has occurred between the neo-X and Y after its initial loss of recombination. This result contrasts with canonical expectations of sex chromosome evolutionary trajectories and could present a new avenue for testing theory relating to sex chromosome evolution - a prospect supported by preliminary simulations of sex chromosome evolution with gene flux.

## Results

### Assembly and annotation of the *G. wheatlandi* genome

We sequenced and assembled the genome of a male *G. wheatlandi*, using a read-partitioning strategy that allowed us to create separate assemblies for the autosomes, the X chromosome, and the Y chromosome (see Methods). This approach resulted in a total assembly length of 519.7 Mb for the autosomes and X chromosome, with 508.5 Mb (97.8%) in 21 chromosome-level scaffolds, matching the expected number of chromosomes from cytogenetic estimates of 2n = 42 (27, 32). Gene annotation for the autosomes and X chromosome, including unplaced scaffolds, yielded 22,010 genes, and *de novo* repeat library and annotation estimated that 25% of the autosomal genome and X chromosome is repetitive, while repeats make up 66% of unplaced scaffolds. BUSCO scores suggest that the completeness of the autosome + X assembly is good (Complete BUSCOs: 97.8% (Single copy: 96.9%, Duplicated: 0.9%), Fragmented: 0.6%, Missing: 1.6%, N = 3640).

To assemble the Y chromosome, we used male-specific k-mers to isolate Y-specific PacBio and Hi-C reads, allowing us to assemble the Y separately from the autosomes and X. The resulting Y assembly was 35.0 Mb long, 33.28 Mb (95.1%) of which was in a single scaffold. This assembly included only the sex-linked region of the Y chromosome, with the 6 Mb pseudoautosomal region (PAR) on Chr12 included in the Chr12 X assembly. Including the Chr12 PAR and unplaced Y scaffolds, we estimate the total length of the Y chromosome in *G. wheatlandi* to be around 42 Mb. Excluding the Chr12 PAR, around 52% of the Y is repetitive, over double that of the autosomes and X chromosomes. Unplaced Y scaffolds have an even higher mean of 75% repeat content. Interestingly, we could find no sign of a PAR on the ancestral Chr19 sex chromosome pair (see SOM results).

### Evolutionary history of the *G. wheatlandi* sex chromosomes

#### Numerous rearrangements on the Y chromosome

Synteny comparisons using 1-to-1 orthologs on X and Y chromosomes revealed extensive rearrangements between them. On the chromosome 19 portion of the Y, the few genes that are left show highly shuffled order relative to the X, implying numerous rearrangements that are too complicated to reconstruct. In contrast, synteny blocks between the Chr12 X and corresponding region of the Y are more contiguous, suggesting fewer rearrangements have occurred in this region. This situation allowed us to infer the most likely order of rearrangements on Chr12. First, the comparison between *G. wheatlandi* and other stickleback species ruled out synteny changes on the X. We did identify a *G. wheatlandi*-specific inversion on Chr12 (Figure S1). However, its breakpoints did not correspond to any inversion breakpoints identified between the X and the Y in *G. wheatlandi*, and no synteny difference is present at this position between Chr12 X and Y (other than Inversion C shown in Figure 1). This inversion thus occurred after the split between *G. wheatlandi* and *G. aculeatus*, but before this region of Chr12 became sex-linked.

**Figure 1.**
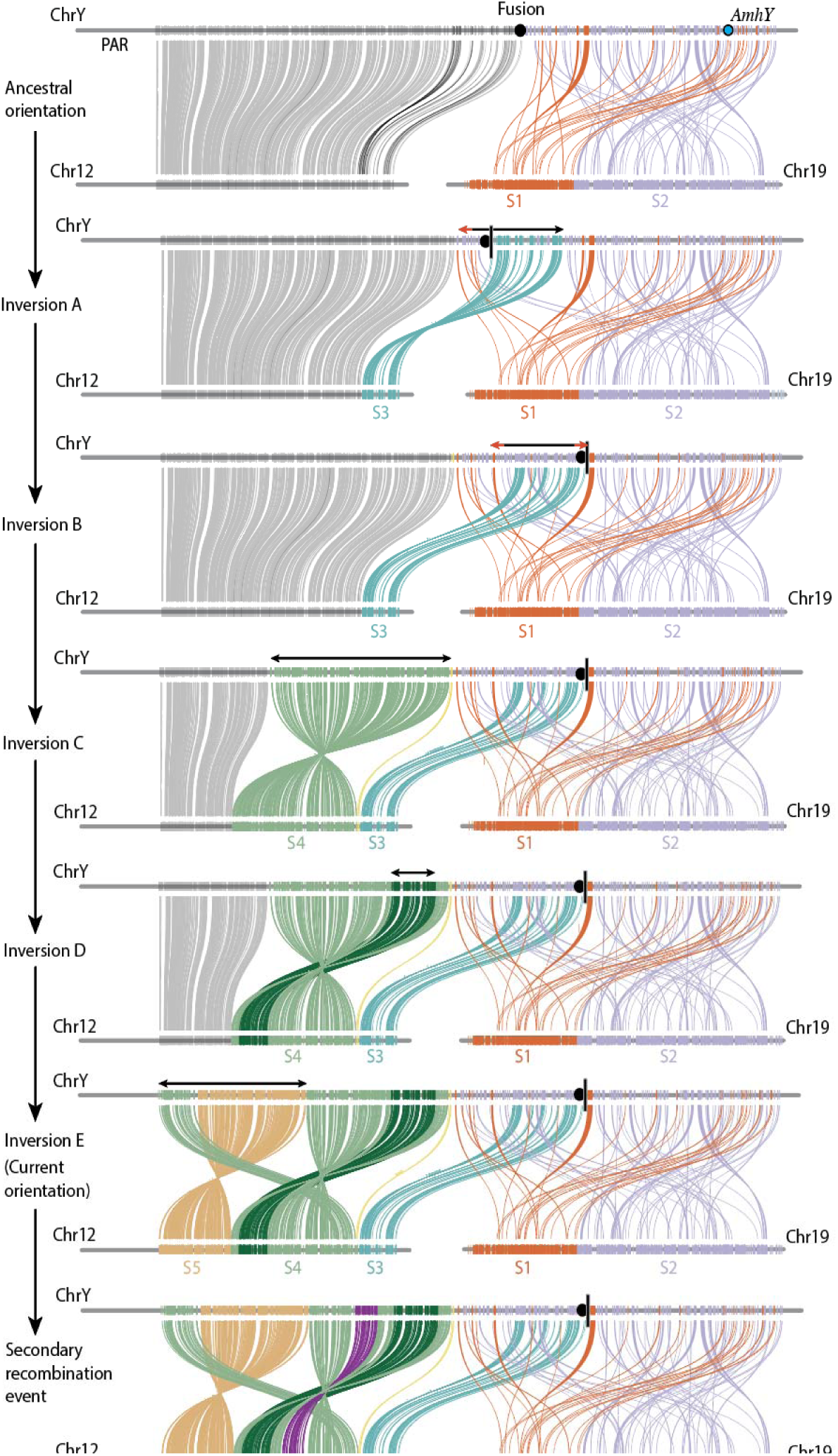
Inferred scenario for rearrangements on the Chr12 Y chromosome, informed by the most parsimonious scenario reconstructed using GRIMM (see methods). Black double-ended arrows above Y chromosomes indicate the approximate extent of the inversion in each step. Red arrow heads indicate lower certainty in the position of the breakpoint. Gene blocks are coloured by strata (inferred by consensus from *F*_ST_, and inversion breakpoints, see Figure 2) except in the case of Inversion D where darker green is used to distinguish the nested inversion within stratum S4. Note that the synteny on Chr19 cannot be reconstructed; the gene orders in this region in the past may not be accurate.

These results provide strong evidence that the X and Y synteny differences are the result of structural changes on the Y chromosome, and that the current X gene order represents the ancestral gene order for the Y prior to sex-linkage. We could thus use a combinatorial approach (GRIMM) to infer the most parsimonious sequence of inversions required to achieve the current Y gene order. This analysis inferred that 5 inversion events occurred on the Y (Figure 1 and see GIF in Appendix 1). Note that the synteny changes that we hypothesise were created by inversions A and B could instead be explained by a single translocation event of this region after the fusion event. However, we have no way of distinguishing between these scenarios; thus, for the sake of simplicity, we invoke only inversions here.

MUMmer (Maximal Unique Match) sequence alignments between inversion breakpoints did not yield substantial evidence for inverted repeats that might imply the mechanism of inversion. However, we cannot rule out that such repeats exist, as these sequences can be small and many of these breakpoints were repetitive and may therefore contain assembly errors.

Finally, we placed the hypothesised location of the fusion involving the ancestral chr19 Y and the neo-Y adjacent to the last gene on the neo-Y (see vertical black line on the Y in Figures 1 and 2). The location of this on the Y assembly is a hypothesis based on the predicted inversions that have likely taken place around the fusion point.

**Figure 2.**
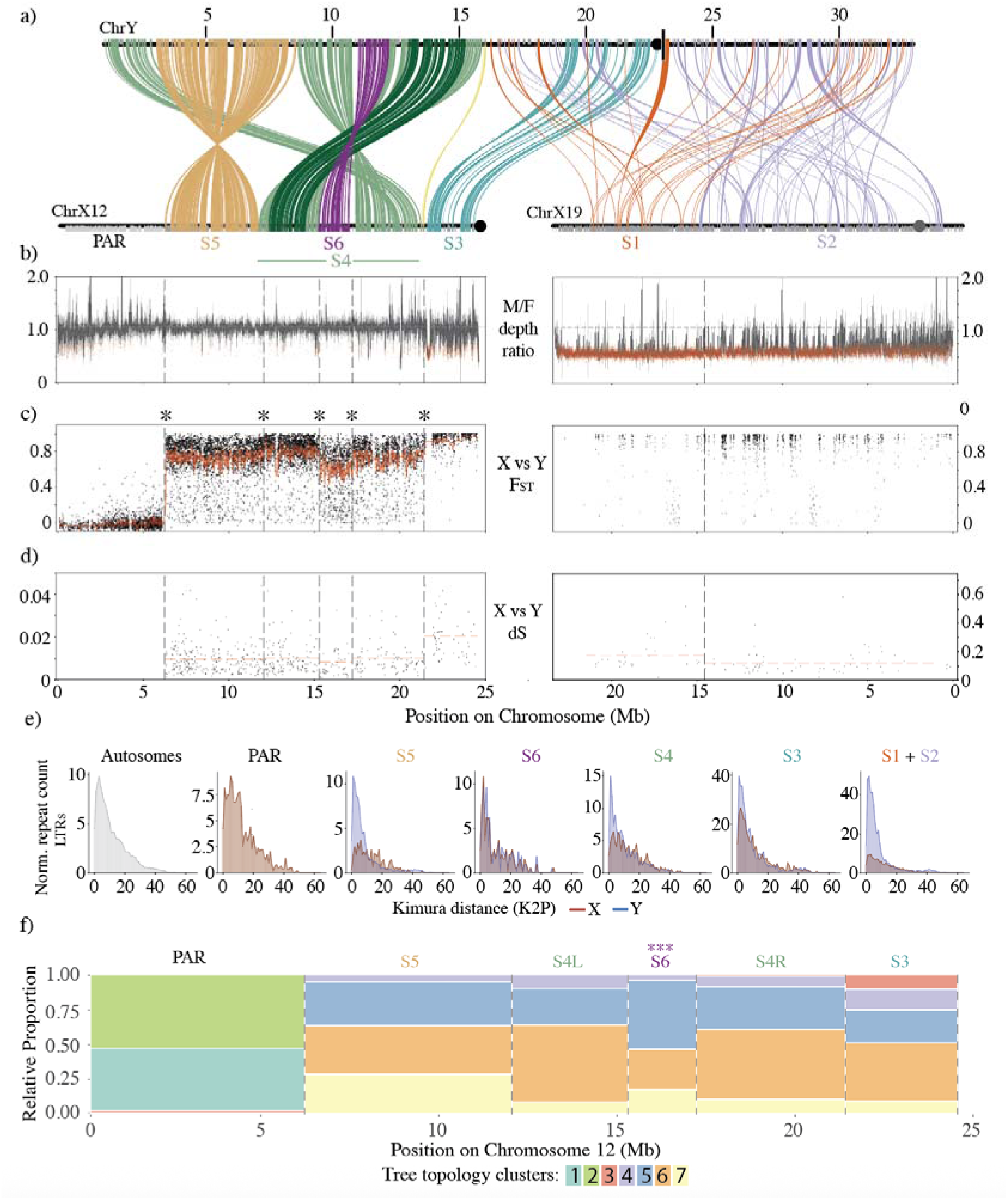
Characterisation of the *G. wheatlandi* sex chromosome. **a)** X vs Y Synteny between 1-to-1 orthologs, with genes and links coloured by inferred stratum. **b)** Male / Female read depth per 1kb window, smoothed over 10 windows. Red points denote sequence windows that have likely been lost from the Y. The dashed horizontal line represents the autosomal male / female read depth average. **c)** Phased X vs Y *F*_ST_ in 1kb windows. The red line is the smoothed average over 10 windows. Dashed vertical lines show the locations of strata boundaries. Asterisks denote those supported by statistically inferred changepoints in the *F*_ST_ distribution. **d)** X vs Y *d*_S_ for 1-to-1 orthologs. Horizontal red dashed lines represent the average *d*_S_ per stratum (defined using the *F*_ST_ changepoint analysis). Note the differing Y axis ranges for Chr12 and Chr19. **e)** LTR age distributions (excluding LTR_2648) for Autosomes, PAR, and X vs Y comparisons for individual strata. **f)** Per-stratum phylogenetic topology proportions. Trees were constructed using phased SNPs in 100kb windows, with a step size of 10kb. Topology cluster colours defined underneath panel f) do not relate to any colours in panel a).

#### Extensive sequence degeneration and gene loss on the Y

We first characterised the extent of sequence loss in the non-recombining region by comparing male and female read depth along the length of the two X chromosomes. In total we found that approximately 19.2 Mb of the chr19 X shows haploid coverage in males and is thus likely missing from the Y chromosome. This constitutes approximately 82% of its original length based on the size of the X chromosome (Figure 2b, S4). In contrast only approximately 1.7 Mb (7%) of the chr12 X was missing on the Y. The loss is restricted mostly to the end of the X chromosome adjacent to the fusion point.

We then used gene annotations to investigate the extent of gene loss on the Y chromosome. Our annotations predicted 1106 and 1072 genes on the Chr12 X and Chr19 X respectively, and 966 genes on the Y chromosome. To estimate the extent of gene loss in different strata, we identified X-genes with intact orthologs on the Y chromosome (Table 1). Gene loss was extensive on the Chr19 region of the Y (S1 & S2), with only 102/1072 (9.5%) X genes having intact homologs. Gene loss was less extreme in the Chr12 region of the Y, with 738/814 (90.7%) of sex-linked X genes retaining an identifiable Y homolog.

**Table 1.** Gene content of strata on the *G. wheatlandi* sex chromosomes. Bottom row.

|  | S1 | S2 | Chr12_PAR | S3 | S4* | S5 | S6 | Total |
| --- | --- | --- | --- | --- | --- | --- | --- | --- |
| **X genes** | 561 | 511 | 292 | 81 | 132, 169 | 344 | 88 | 2178 |
| **Retained Y homologs** | 72 | 30 | n/a | 65 | 116, 153 | 322 | 82 | 897** |
| **% retained** | 12.48% | 5.87% |  | 80.25% | 84.09%, 89.94% | 90.70% | 89.77% | 41.18% |

\* The two numbers for each row represent the two sides (left, right) of S4 which are separated by S6.
\*\*15 Y genes have no detectable homolog. 54 Y genes could not be processed with Orthofinder. Total Y genes = 966.

#### Synteny and sequence differentiation distinguish multiple evolutionary strata

As sequence degeneration and rearrangements on the Chr19 part of the Y chromosome are so extensive, it was not possible to infer strata based on synteny or metrics of sequence differentiation in this region. However, *G. wheatlandi* and *G. aculeatus* share the oldest stratum on Chr19 (S1) (30). We therefore defined regions belonging to this stratum on the *G. wheatlandi* X chromosome based on orthology with genes in *G. aculeatus*, which has a clear delineation between the oldest and younger strata (21). We were unable to determine whether the rest of chromosome 19 lost recombination in one mutational step, or several. Thus, while we assign this region to a single stratum (S2) for simplicity, we cannot rule out that additional strata once existed.

For the Chr12 X and the corresponding section of the Y, we inferred strata as regions at which changes in X vs Y *F*_ST_ aligned with structural differences between the X and Y. Changepoint analyses of *F*_ST_ identified three changepoints that aligned to the breakpoints of three major inversions between the Chr12 X and the Y (Figure 2a,c). Male / female read depth ratio on the neo sex chromosome was lowest closest to the fusion point, which was also the region with maximal X vs Y *F*_ST_ and *d*_S_. This region also corresponds to an inversion between the X and Y (Inversion A, Figure 2), the left-most breakpoint of which aligns to an *F*_ST_ distribution changepoint. We therefore inferred that this region (S3) is the oldest stratum on the neo sex chromosome. Read depth ratio along the rest of the neo sex chromosome deviated little from the autosomal average, concordant with the results of our genome annotation showing limited Y gene loss in this region. However, two additional *F*_ST_ distribution changepoints aligned with the breakpoints of Inversions C and E, allowing us to infer strata S4 and S5.

Intriguingly, we identified two more *F*_ST_ changepoints indicative of an additional stratum on the neo-Y (S6) with an obvious 2 Mb depression in *F*_ST_ is seen at 14.6 - 16.6 Mb (stratum S6, Figure 2c). However, the changepoints flanking S6 did not align to any inversion breakpoints. Further, the lower *F*_ST_ observed in S6 disrupts the expected stepwise pattern of strata formation on sex chromosomes. When non-recombining regions of sex chromosomes expand, theory predicts that they do so via progressive mutations that either encompass or overlap with other previously sex-linked regions (e.g. overlapping inversions like Inversion C and Inversion E, Figure 1). Age of strata, and consequently X vs Y differentiation, should therefore decrease from oldest to youngest in a stepwise manner via adjacent strata. This expectation holds for chromosome 19 (the oldest region of the sex chromosome) relative to S3 (the oldest region of the neo-Y). The next region to become sex linked spanned 11.74 - 21.29 Mb and was created by Inversion C (Figure 1). As this region would have become sex-linked simultaneously, it should have similar X vs Y differentiation across its length, thus, the depressed *F*_ST_ observed in the centre of this region was surprising.

#### Evidence for secondary recombination within an existing stratum

The non-stepwise pattern of *F*_ST_ identified within S4, and specifically the low *F*_ST_ in region S6 suggests that this region has been undergoing X/Y differentiation for less time than its flanking regions. This, in turn, implies recombination in this region has occurred after the initial loss of recombination in stratum S4, despite the loss of homology resulting from Inversion C (Figure 1). We assessed support for this hypothesis from three other measures of X/Y differentiation.

First, *d*_S_ estimates at four-fold degenerate sites between 1-to-1 homologous genes on the X and Y showed the same pattern as *F*_ST_, with a depression of sequence divergence within S6 relative to the rest of S4 (Figure 2d). Elsewhere on the sex chromosomes, estimates of *d*_S_ were consistent with our hypothesis for the order of strata formation.

Second, we estimated the age distribution of repeats in each stratum on the X and Y. Repeats are known to accumulate in non-recombining regions of sex-limited chromosomes (33–35). We thus expect a burst of repeat insertions on the Y to coincide with the formation of a new stratum. If S6 truly has recombined more recently than the rest of S4, the burst of new repeat insertions on the Y should be less obvious in this region. We found that all strata on Chr12 showed a clear enrichment of young repeats on the Y chromosome relative to the same region on the X. Surprisingly, Stratum S6 showed a Y-specific burst of old repeats; however, further investigation revealed that this pattern was driven by a single type of repeat, an unclassified LTR (LTR_2648, See SOM Results and Figure S2 for details). After accounting for this single anomalous repeat, we see no other signal of increased numbers of young repeats on the Y in this stratum for any other repeat family (Figure 2e), consistent with our hypothesis that this stratum has been sex-linked for less time than its flanking regions (Figure S2). In contrast, when we removed this repeat from our analyses in other strata, we still observe a burst of young repeats on the Y (Figure 2e), owing to the concentration of LTR_2648 copies in just a single location within S6 on the Y.

Third, phylogenetic analyses based on phased X and Y SNP calls showed that S6 has a distinct phylogenetic signal (Figure 2f) from S4. Tree distances in S6 were significantly elevated in comparison to the S4 flanking regions based on permutation tests (p < 0.001, n=1000, Figure S3). In particular, topologies in which X and Y sequences are reciprocally monophyletic (topology cluster 6 Figure S3c) were less frequent relative to topologies where Y sequences were monophyletic but nested within the X sequence clade (topology cluster 5). As reciprocal monophyly is indicative of more complete lineage sorting, its relative absence in S6 is also consistent with more recent recombination in this region relative to S4.

In summary, we find evidence in support of a secondary recombination event in S6 following the initial loss of recombination in S4 from four analyses based on two independent datasets: X vs Y *F*_ST_ and phylogenetic tree topologies calculated from phased population sequencing data, and *d*_S_ and repeat ages, based on the phased X and Y chromosome assemblies.

#### Signatures of gene flux on sex chromosomes

The evidence for a double-crossover event between the X and Y in their “non-recombining” region suggests an interesting possibility. Gene flux caused by double crossovers and gene conversion could produce signatures of the evolutionary forces acting on sex chromosomes. These signatures might be used to test hypotheses for what maintains recombination suppression between X and Y chromosomes: sexually antagonistic selection, or constraints on the production of appropriate structural changes. In the following, we use “gene flux” to refer to recombination resulting from double crossovers as distinct from recombination involving only a single crossover between the X and Y.

To investigate this idea, we simulated sex chromosomes containing a PSDL which evolves with gene flux and either with sexually antagonistic loci (SAL) or without them (see Methods for details). Results showed that the relative rates of mutation and gene flux determine whether useful signals can be extracted from sex chromosome sequences. When the relative rate of gene flux is very low, mutation and drift erase the signatures of gene flux events (Figure 3a, b). At the other extreme, when the relative rate of flux is high, it homogenizes the X and Y, and again no signal is detectable (Figure 3e, f). However, when rates of mutation and gene flux are comparable, the cumulative effects of double crossover events over time yield patterns that can be used to identify loci resisting recombination (Figure 3c, d).

**Figure 3.**
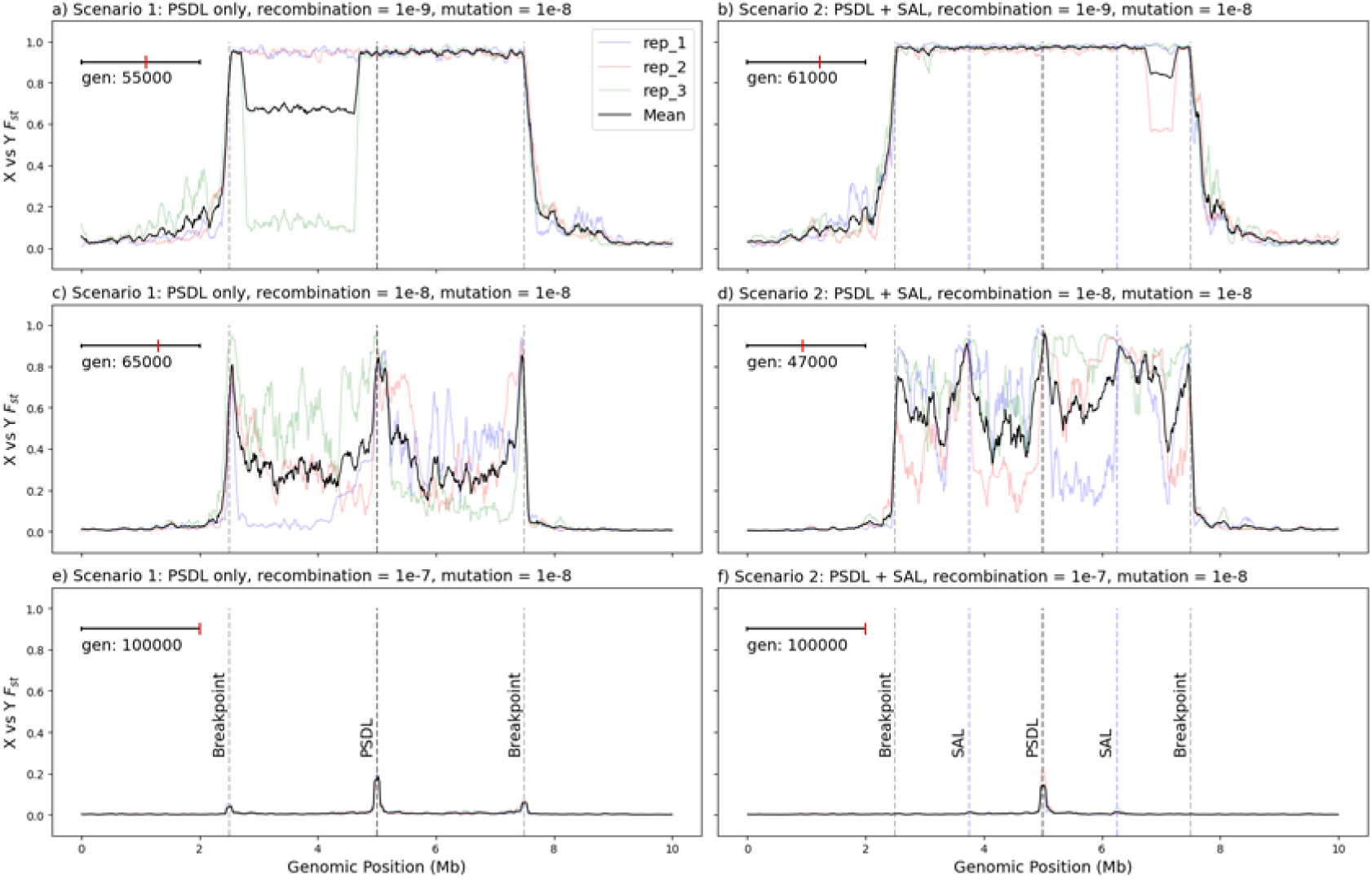
Simulations of sequence differentiation on an evolving sex chromosome with a 5Mb non-recombining region denoted by “Breakpoints” and with gene flux possible within this region via double crossovers between X and Y. The simulated non-recombining region contained either a PSDL only (left panels), or PSDL + Sexually Antagonistic Loci (SAL) (right panels), and simulations were conducted for three combinations of baseline mutation and recombination rates (rows). X vs Y *F*_ST_ was calculated in 1kb windows, and the rolling Mean *F*_ST_ was calculated using 10 windows with a 1 window step. Plotted generations were chosen to be indicative of results highlighted in the text. See Appendix 2 for an animated version of the simulations.

Consider a double crossover event which occurs within the sex chromosome inversion. If this region does not contain a breakpoint, the PSDL or a SAL the X and Y recombinants are free to fix, thus decreasing divergence between the X and Y (Figure 3c). In contrast, consider the outcome when the double crossover event spans a region that contains one such locus. In these cases, recombinant haplotypes are selected against. Ongoing divergence thus produces peaks of *F*_ST_ around these sites (Figure 3d). This pattern establishes within 10^4^ generations after the origin of the inversion (Appendix 2).

While these patterns are easily noticeable when averaging the results of three replicates, single replicates often show transient *F*_ST_ noise driven by drift that creates spurious peaks and masks the true PSDL and breakpoint signals (Figure 3c, d, Appendix 2). However, in both the PSDL-only and PSDL+SAL scenarios, the *F*_ST_ at PSDL, SAL (if present), and breakpoints remained consistently strong in all replicates throughout the simulations (Appendix 2). These simulations therefore suggest that estimating patterns of past gene flux in sex-linked inversions could indeed allow us to identify candidate regions containing a PSDL or SAL (*F*_ST_ peaks), as well as regions where such loci are absent, or, in the case of SAL, weakly selected for.

## Discussion

Phased sex chromosome assemblies enable the identification of structural rearrangements between gametologs. Generating phased assemblies of the *G. wheatlandi* sex chromosomes has allowed us to associate evolutionary signatures indicative of strata (e.g. plateaus of high X vs Y differentiation) to the proximate mechanisms that created them (i.e. inversions), greatly increasing confidence in proposed strata locations, age, and content. With this information in hand, we were able to observe several interesting phenomena on the sex chromosomes of *G. wheatlandi*. We identified a large number of inversions, some of which likely increased the length of the non-recombining region of the sex chromosome, while others likely had no effect on recombination, but fixed regardless. We also identified the likely location of the surviving centromere on the Y chromosome and observed an interesting case of a Y-specific repeat expansion. We were unable to find one of the two expected PARs, despite expectations that it is necessary for proper sex chromosome segregation. While not directly relevant to the central result of this study, further discussion of these interesting observations can be found in the supplementary materials section “On the biology of *G. wheatlandi* sex chromosomes”.

### Recombination within non-recombining sex chromosomes

The most intriguing finding of this study is the strong evidence that recombination has occurred within an existing stratum after its initial loss of recombination, likely via double-crossover mediated recombination between inverted haplotypes. While double-crossover recombination events between autosomal inversion haplotypes have been observed in fruit flies (36, 37), butterflies (38), fish (39), and plants (40), our results represent the first observation of this phenomenon in sex chromosomes. We know of only one other empirical example of recombination occurring on a sex chromosome after recombination has initially been lost, which was in the Spanish marbled white butterfly, *Melanargia ines* (41). However, recombination loss in this case is due to achiasmatic female meiosis, and the distal and stepwise nature of the proposed recombination events in this species suggest that they were most likely single crossover events. Thus, it is likely that the mechanism of ZW recombination in *M. ines* is different from the double-crossover mediated recombination event proposed in the present study.

One other interesting observation from our results is the almost-perfect association of the boundaries of the double crossover event that created stratum S6 with the breakpoints of an observed inversion between *G. wheatlandi* and its sister species *G. aculeatus* in the same region. This may be a coincidence. However, it may reflect the presence of one or more fragile sites in this region, i.e. loci that are repeatedly involved in double strand breaks leading to recombination or structural rearrangements (42). There are in fact several rearrangements in this region of Chr12 between multiple stickleback species (Figure S1), and the enrichment of tandem repeats near to the S6 boundaries (see SOM results and Figure S2) is consistent with expectations for fragile sites. It is therefore possible that both the interspecific inversion and the secondary recombination event that created stratum S6 were both facilitated by the fragility of this tandem repeat-rich genome region.

### Gene flux on sex chromosomes as a tool for testing models for their evolution

A persistent mystery in the field of sex chromosome evolution is why recombination loss is maintained in the long term when it comes with the heavy cost of deleterious mutation accumulation (43). Two possible explanations are either that mutations required to restore recombination (e.g. reversions) are rare (25), or that selection favours the non-recombining haplotype due to some advantage conferred by the linkage between the PSDL and one or more loci within the non-recombining region (16, 17, 19). Debate is ongoing as to which of these processes is more important in nature, but only with empirical data can we hope to definitively answer this question.

Unfortunately, empirical tests of both the constraint theory, and the adaptive linkage theory present several challenges. From an experimental perspective, inversion/reversion mutations occur over time scales that likely make experimental tests untenable. CRISPR offers exciting possibilities for inducing reversions (44), though to our knowledge this has not yet been tried in a sex chromosome context and may not be possible in all taxa. Comparative genomics between closely related species with shared sex chromosomes could potentially be used to look for signals of reversions. However, this would rely on the lucky distribution of polymorphisms on a well sampled phylogeny to infer that a reverted orientation is derived.

The occurrence of gene flux between the X and Y, as observed here, could offer much needed empirical perspectives on these theories as they make opposing predictions about the fate of recombinant Y haplotypes. Gene flux is expected to create a so-called ‘suspension bridge’ pattern of differentiation between haplotypes (45, 46), due to the lower rates of double crossovers and gene conversion events in regions nearer to inversion breakpoints. Such patterns have been observed in autosomal inversions of *Anopheles gambiae* (47) and *Drosophila melanogaster* (48). Though conceived in the context of autosomal inversions, this theory can easily be carried over to sex-linked inversions. Under a pure constraint model, double-crossover recombinant Y haplotypes would be free to fix via drift, or via positive selection due to their lower mutation loads, thus creating a suspension-bridge pattern. In contrast, under an adaptive-linkage model, recombination between X and Y haplotypes would be selected against if it breaks linkage between the PSDL and a Y allele that is adaptive when linked to it. Sexually antagonistic loci would thus resist gene flow between haplotypes (49, 50), which would result in localised peaks of differentiation at such loci within the background suspension-bridge pattern (46, 51). Patterns of gene flux between sex chromosomes could thus offer a way to empirically distinguish between the elusive evolutionary processes at work within sex-linked regions.

Our simulations of sex chromosome evolution with gene flux performed here support these expectations. They show that when selection disfavors recombinant X or Y haplotypes, as with sexually antagonistic selection, differentiation persists at selected sites and breakpoints while flux still occurs in the intervals between them. These results are also fully consistent with previous analytic theory (Kirkpatrick and Guerrero 2014) and simulations (Dagilis et al. 2022) of sex chromosome evolution with and without sexually antagonistic selection. While the earlier models assume recombination between the X and Y results from single crossovers, the double-crossover model used here (and possibly gene conversion) do not alter the qualitative outcomes.

In addition, there are several characteristics of sex-linked inversions that may make them better suited for observing patterns of past gene flux than their autosomal counterparts. Most importantly, the enforced heterozygosity of sex chromosomes means that the inversion haplotypes therein likely spend more time in heterozygotes than inversions on autosomes. As gene flux happens only in heterozygotes, this significantly increases the potential for gene flux relative to autosomal contexts. Second, sex chromosomes often harbour large inversions (e.g. >5Mb) (21, 23, 52, 53), which are better suited to identifying spatial patterns in haplotype differentiation. Third, lower effective population sizes, strong linked-selection and sex-biased mutation rates experienced by sex-linked chromosomes likely increase the rate at which young inversion haplotypes differentiate. Finally, the tendency of sex chromosomes to accumulate deleterious mutations raises the likelihood that, when recombination occurs in sex-linked regions that do not contain a breakpoint, PSDL or sexually antagonistic locus, the recombinant Y fixes, and the recombinant X is lost. This would increase X/Y homogenisation in “troughs” and could accentuate peaks at loci resisting recombination. All the above factors should be explored in future simulations to best identify situations in which we might empirically observe the processes discussed above.

With this in mind, how might we interpret the empirical results in *G. wheatlandi* stickleback presented here? First, *G. wheatlandi* is a good candidate system in which the history of gene flux could be evaluated. Estimated mutation and recombination rates from the closely related *G. aculeatus* are of a similar order of magnitude (54, 55). Inversions are large (∼3-10 Mb), and strata ages vary by an order of magnitude. Second, the recombination event identified here in a sex-linked inversion tells us that such a mechanism is possible in this system. Third, we do see signs of a suspension bridge pattern of differentiation in stratum S5 (Figure 2), which could indicate a lack of sexually antagonistic selection in this region. However, this dataset suffers from a relatively low sample number, making *F*_ST_ estimates noisy. In addition, having data from multiple populations would be somewhat akin to the multiple replicates in our simulations, and could allow us to find high *F*_ST_ peaks shared across populations. Adding data from additional samples and populations in this species would thus greatly improve our ability to relate empirical results in this system to theoretical expectations.

In conclusion, our results from *G. wheatlandi* herein highlight gene flux as an important mechanism to consider when studying the evolution of sex chromosomes. Indeed, it would be interesting to incorporate this process into theoretical models of sex chromosome evolution (e.g. 17, 25, 56). Further, this observation has inspired a potentially powerful empirical framework, which our simulations suggest could help us infer the evolutionary forces acting within non-recombining sex-linked regions.

## Materials and Methods

### Sample collections

The *G. wheatlandi* specimen used for genome assembly was a laboratory-reared adult male resulting from a cross between wild fish collected from Canal Lake (44.49830, −63.90205) in Nova Scotia, Canada by Anne Dalziel. Two male siblings from the same cross were used for Hi-C sequencing to scaffold the PacBio assembly (see below). The samples used for population genetics analyses were comprised of: four males and four females collected from Rainbow Haven Beach (44.654857, −63.42113) in Nova Scotia, Canada by Anne Dalziel in 2018; three males and three females collected from Demarest Lloyd State Park in Dartmouth, Massachusetts, USA by CLP in 2007; two males collected from Wells, Maine, USA by Bill Rowland in 2003; and two males and two females collected from Baie de L’Isle-Verte National Wildlife Area in Quebec, Canada by Louis Bernatchez in 2003. All experiments involving animals at the University of Bern were approved by the Veterinary Service of the Department of Agriculture and Nature of the Canton of Bern (VTHa# BE4/16, BE17/17 and BE127/17).

### DNA and RNA extraction and sequencing

For the genome assembly, high-molecular-weight DNA was extracted from the blood of a single *G. wheatlandi* male following the protocol described by (21). The extracted DNA was subsequently used to construct a SMRTcell Express library for sequencing on a PacBio Sequel II at the University of Bern Next Generation Sequencing Platform. Liver tissue from the same individual, along with that from his two siblings, was used to prepare a Hi-C sequencing library. The library construction was performed using the Phase Genomics Proximo Hi-C Animal Kit (Phase Genomics, Seattle, WA) and Hi-C libraries were sequenced for 300 cycles on an Illumina NovaSeq SP flow cell at the University of Bern Next Generation Sequencing Platform. To guide gene annotation, we used previously published RNA seq data (57), which can be found at the accession PRJNA746773.

For population genetics analyses of the sex chromosome, DNA from eight wild-caught *G. wheatlandi* individuals collected at Rainbow Haven Beach, Nova Scotia, Canada was extracted using the phenol–chloroform method. The extracted DNA was used to construct Illumina TruSeq DNA libraries, which were sequenced for 300 cycles on an Illumina NovaSeq S1 flow cell at the University of Bern Next Generation Sequencing Platform. DNA from the samples collected from Massachusetts, Maine, and Quebec was extracted from fin clips in ethanol with the Qiagen DNeasy kit, and Illumina TruSeq DNA libraries were prepped and sequenced at UT Austin GSAF on a HiSeq 4000.

In this study, we also used available whole-genome sequencing data of 15 males of *G. wheatlandi*, 15 females of *G. aculeatus*, and 15 interspecies crosses of *G. wheatlandi* (paternal) and *G. aculeatus* (maternal) from (30).

### *De Novo* genome assembly and phased sex chromosome assembly via kmer-based read X and Y read partitioning and

To minimize the interference of Y chromosome reads with autosomal assembly and to achieve a complete assembly of the Y chromosome, we classified the raw PacBio and Hi-C reads into two groups: 1) Autosomes + X chromosome, 2) the Y chromosome only. We did this by first identifying male-specific kmers (MSK) by comparing kmers present in the whole-genome sequencing data of the four males and four females described above. This was done using the software SRY (v1.6) (58). SRY assigns reads into groups based on the density of MSKs that align to them. Given their short length, Hi-C reads were assigned to the Y read group if at least one MSK aligned. For PacBio reads, a threshold density of MSKs per read was used. As some regions of the *G. wheatlandi* sex chromosomes are not strongly differentiated (30), the density of Y specific kmers in some regions may be low. We thus first produced a preliminary assembly for the autosomes, X and Y chromosomes (see below for assembly methods). We then reran read assignments in SRY multiple times, testing different per-read MSK density thresholds from 2.0 to 4.0 with a step size of 0.2. We aligned each of the resulting Y-specific readsets to our preliminary genome assembly using minimap 2 (v2.28) (59) and calculated the sequencing depth with a window size of 1kb using Deeptools (v3.5.5) (60). Lower MSK values resulted in higher Y-read depth as fewer MSKs were required per read to define it as Y specific. We chose a final MSK value of 2.4, as this was the point at which the increase in Y-read depth with decreasing MSK plateaued. Using this threshold, the assembly pipeline was re-executed to generate two separate assemblies: one for the autosomes and X chromosome, and another for the Y chromosome.

For all genome assemblies above, our pipeline was as follows. The two readsets (autosomes + X readset and Y readset) were independently assembled at the contig level using Flye (v2.9.5) (61) with default parameters. The same long-read data were used for polishing the primary assemblies by racon (v1.5.0) (62) with default parameters separately, followed by the polish with corresponding Hi-C data with pilon (63). Duplicates in assemblies were purged by Purge_Dups (64). Hi-C reads from two datasets were aligned against the contig assemblies by Chromap (v0.2.7) (65) and scaffolded by YaHS (v1.2.2) (66). Finally, gaps were closed with the help of long reads conducted by TGS-GapCloser (v1.2.1) (67).

The quality of the final autosomal and X assemblies was evaluated by BUSCO (v5.8.0) (68). Unfortunately, it is not possible to reliably assess the completeness of our Y assembly via BUSCO genes as those that are missing may either be unannotated, or have been lost via degeneration of the Y chromosome.

### Genome Annotation

The autosomal, X and Y assemblies were merged during the annotation step. A two-step pipeline was used to annotate the genome assembly. First, a repeat library was constructed using EDTA (v2.2.0) (69), and then unknown repeat elements were further classified with DeepTE (70). RepeatMasker (v4.1.3) (71) were used to generate a final annotation of repeats.

Next, RNA-sequencing data from eight *G. wheatlandi* individuals (four males, four females) and described above were used to aid in genome annotation. Raw sequences were downloaded and mapped against the soft-masked genome assembly by STAR (v2.7.11b) (72) with the two-pass mode. Genome annotation was done by Braker (v3.0.8) (73) with the integration of RNA and external protein database of Actinopterygii from NCBI (taxid:7898). Lastly, the functional annotation was conducted by eggnog-mapper (v2) (74).

### Genomic Synteny Analyses between X and Y chromosomes

First, whole genome comparison was conducted using MUMmer4 and NUCMER (75) between *G. wheatlandi* and *G. aculeatus* to confirm the sex chromosomes. Next, nonredundant coding sequences were used for synteny analysis. Longest isoforms of each gene were kept using AGAT (76). We then used MCScan (77) in JCVI package to compare synteny at gene level between X and Y chromosomes.

#### Inference of inversion orders

To infer the order of structural rearrangements on the Neo-Y chromosome, we first established the ancestral gene order on the X, i.e. after the split between *G. wheatlandi* and *G. aculeatus*, but prior to Chr12 becoming sex linked via fusion to the Chr19 Y. We did this via gene order comparison between *G. wheatlandi* chr12 X and the orthologous chromosomes of three other stickleback species for which genome assemblies exist, *G. aculeatus*, *Apeltes quadracus*, and *Aulorhynchus flavidus* using the same methodology as described for comparisons of the X and Y chromosomes above.

We then inferred the most parsimonious (i.e. fewest) sequence of inversions that transform the ancestral chr12 X gene order into the present Y chromosome gene order. At its core, this is a combinatorial mathematics problem akin to the pancake flipping problem (78). To solve this problem, we used GRIMM v2.01 (79) implemented in the R package GRSR (80). One drawback of this approach is that GRIMM does not require successive inversions to overlap with each other, which is a requirement of progressive recombination loss on sex chromosomes. We thus broke the problem down into several sub tasks focusing on specific regions of the X and Y chromosome. This was guided by the *F*_ST_ results which gives a broad indication of the order in which each region lost recombination. Combining the outputs from GRIMM from these subtasks, specifically strata sets (S1 + S2 + S3) and (S4 + S5), allowed us to put a complete scenario together for rearrangements between chr12 X and the Y.

### Whole-genome alignment and variant calling

To identify genomic variants, we combined three datasets into our analysis. Raw reads were first quality checked by fastqc (81) and then trimmed by Trimmomatic v0.38 (82). Trimmed reads were mapped to the reference assembly with Y chromosome removed using bwa (v0.7.11) (83) and default parameters. All bam files were then sorted with Samtools (v1.20) (84) and duplicates were removed by sambamba (85). Variant calling was done using HaplotypeCaller, and joint genotyping was run by combining all individuals for the population with GATK4 (v4.6.1.0) (86), followed by hard filtering according to the GATK best practice pipeline.

For variant calling, the samples were separated into three subsets based on the design and sequencing quality: (1) 15 pedigrees between females from *G.aculeatus* and males from *G.wheatlandi*, from (30); (2) Eight samples from Rainbow Haven Beach, Nova Scotia, Canada; (3) twelve samples from Maine, USA (N=2), Massachusetts, USA (N=6) and Quebec, Canada (N=4). For dataset 1, we used VCFtools (0.1.16) (87) to retain SNPs with a minimum quality score above 30, a genotype quality above 20, and a genotype depth of at least 8. To reduce bias from extreme sequencing depths, we further filtered out sites where the population mean depth coverage was below 15× or above 75×. Given that the data originated from inter-species crosses, we followed the phasing protocol described in. Specifically, for each heterozygous SNP in the offspring, we used parental genotypes to determine the paternal and maternal alleles. Paternally inherited alleles were transmitted via sperm carrying a Y chromosome (in sons) or an X chromosome (in daughters), allowing us to independently sample 15 *G. wheatlandi* X chromosomes and 15 *G. wheatlandi* Y chromosomes from the wild. After phasing, we refined the SNP matrix by removing sites with fewer than five individuals in each sex. For dataset 2, we applied the same filtering criteria for SNP quality, genotype quality, and depth but adjusted the sequencing depth thresholds, retaining sites with a population mean depth coverage between 10× and 50×. For dataset 3, due to lower sequencing depth, we strictly retained only SNPs meeting the minimum thresholds of a quality score above 30, a genotype quality above 20, and a genotype depth of at least 8, without additional depth filtering.

Finally, we produced two variant callsets based on the analyses above: (A) a callset of phased haploid genotypes of the offspring from the inter-species cross in dataset 1; (B) a matrix including paternal individuals from dataset 1, all individuals from dataset 2 and 3.

### Characterising evolutionary history and strata on the sex chromosomes

First, we estimated the extent of sequence loss on the Y chromosome using the filtered BAM files to calculate average normalised (RPKM) read depth for males and females in 1kb windows with deepTools2 (60). We then plotted the histogram for (mean Male / mean Female) normalised read depth per 1kb window to identify thresholds to assign windows as being X specific (i.e. lost from the Y) (Figure S4). Note that it is possible for only a portion of a single 1kb window to be X-specific, and, while based on the histogram, the choice of thresholds to assign X-specific windows is still somewhat arbitrary. However, despite these caveats we believe this approach still gives a relatively reliable estimate for the proportion of Y sequence that has been lost.

For sequence windows present on both X and Y, differentiation was assessed using *F*_ST_, which was calculated using Vcftools (v0.1.16) again for 1kb windows. We used the resulting *F*_ST_ values to identify putative evolutionary strata. Evolutionary strata on sex chromosomes are regions of similar X vs Y differentiation which result from the same recombination-loss event. To identify stratum boundaries, we used statistical changepoint analysis of *F*_ST_ distributions with the Python package “ruptures” (88).

We then calculated divergence between genes present on both X and Y chromosomes. We used Orthofinder (v2.5.5) (89) to identify orthology and parology between sex chromosome genes. As input, we extracted protein sequences corresponding to the longest isoforms from the X and Y chromosomes. To enhance the accuracy of orthologue detection, we incorporated additional outgroup species, including *Gasterosteus aculeatus* (both autosomal and Y chromosome assemblies) (21), *Apeltes quadracus* (autosomal assembly only) (28), and *Pungitius pungitius* (autosomal assembly only) (90). The species tree used for phylogenetic inference was obtained from (57). Ortholog identification was conducted using the ultra-sensitive mode to optimize detection sensitivity. As the final dataset, we selected “phylogenetic hierarchical orthogroups” that included representatives from all species, ensuring a comprehensive comparative framework for assessing orthology.

Next, 1-to-1 orthologs between X and Y chromosomes from *G. wheatlandi* were extracted and protein sequences of each pair were aligned by PRANK (91). Then, pal2nal (92) was used to align the sequences of coding regions under the guide of protein alignments. In the end, KaKs_Calculator 3.0 (93) was adopted to calculate *d_N_* and *d*_S_ value of each gene pair. For *d_N_*/*d*_S_ values, genes with a value of 99 were not included in the downstream analysis. Stratum boundaries from the *F*_ST_ changepoint analyses were used to calculate mean per-stratum *d*_S_ values.

To examine evolutionary history on chromosome 12 in more detail we used window-based phylogenetic analysis, capitalising on the phased X and Y SNP calls from Sardell et al., (30) data. SNP calls were masked for any remaining repeat rich regions and then converted into consensus haplotypes for chromosome 12 using bcftools consensus (94). Phylogenetic trees were constructed for 100kb windows with a step size of 10kb along the chromosome using the Automated-Window-Sliding pipeline (https://github.com/ggruber193/automated-window-sliding/, accessed on 18th May 2026). Trees were made using the IQ-tree algorithm.

Phylogenetic trees were analysed using the R package treespace v1.1.4.4. (95). Trees were clustered according to their pairwise similarity defined by a distance metric that compares the time to the most recent common ancestor for each pair of tips (96). Tree distances were visualised using PCA (calculated by treespace) and topology clusters were identified using the findGroves function using the Ward.D clustering algorithm. Topologies closest to the cluster median were used to represent each cluster and the cluster memberships of topologies were plotted along the genome. Stratum boundaries from the *F*_ST_ changepoint analyses were used to calculate partition and compare topology proportions across strata (see below).

To assess whether the genomic window S6 (coordinates 15.2965–17.2135 Mb) is phylogenetically distinct compared to the background of S4, we implemented a paired-permutation test designed to control for reference-set size and local variance. To create the null distribution, for each permutation, a random 1.917 Mb window was sampled from stratum S4 (comprising S4L: 12.0455–15.2965 Mb and S4R: 17.2135–21.4005 Mb). A temporary “Reference Set” was dynamically defined for each iteration as the remainder of the background trees after excluding that iteration’s sampled window. For each permutation we calculated the median of the pairwise tree distances between the randomly-sampled window and the background reference of each permutation. We also calculated the median distance from S6 trees to the reference background of each permutation to produce the “observed distribution”. The final p-value was defined as the proportion of permutations in the null distribution for which the median tree distance between the randomly-sampled window and the background S4 region exceeded the median of the observed distribution.

### Repeat content on the sex chromosomes

To further investigate the divergence between X and Y chromosomes, we performed a repeat enrichment analysis. A de novo TE library was extracted from the output of EDTA (v2.2.0) (69), and further classified with DeepTE (70). Whole-genome TE annotation was subsequently performed with RepeatMasker (v4.1.3) (71) and the Kimura 2-Parameter (K2P) distance was calculated for each repeat element. A custom script was used to extract K2P values from the RepeatMasker output. To avoid redundant calculations of the same genomic fragment, only the primary domain with the highest score was retained in cases of overlapping matches. The identified repeats were classified into four distinct categories (LTR, non-LTR, TIR and non-TIR) and further grouped based on strata information. Finally, the data were smoothed using the locally estimated scatterplot smoothing (LOESS) model implemented in the Python package “statsmodels” (97).

### Gene flux simulations

We used SLiM v5.2 (98) to run forward genetic simulations of sex chromosome evolution. The base simulation setup was a population of 1000 individuals with two 10Mb genomes simulated under a Wright-Fisher model (99). Each generation we forced all individuals to be XY. We defined the Y chromosome by an introduction of a neutral locus (PSDL) at 5MB in the genome, whereas the X chromosome was defined by PSDL absence. We simulated two main scenarios: 1) PSDL capture by the inversion, and 2) PSDL and two sexually antagonistic loci captured by the inversion. In the first scenario we let neutral mutations accumulate for 10000 generations before introducing a 5Mb inversion centered on the genome in a single Y chromosome. Here, selection (*s* = 0.1) acts only on the breakpoints with an equal, but opposite fitness effect (*w* = *w* × (1.0 + *s*)) in X and Y, such that the inversion is beneficial on the Y chromosome and deleterious on the X chromosome. In scenario two, we additionally initiated two sexually antagonistic loci (SAL) in a Y chromosome of a single individual at positions 3.75Mb and 6.25 Mb in the first generation. Here, selection favours one allele at the SAL on the Y chromosome and selects against recombination events that moves it onto the X (and *vice versa* for the other allele). At generation 10000 we introduced a neutral, 5Mb inversion centered on the genome in a single Y chromosome, capturing the PSDL and SAL. In both scenarios, we allowed only an even number of crossovers to occur inside the inversion. If the inversion was lost from the population, we restarted simulation at generation 10000. We simulated three recombination rates (*r* = 1×10^−9^,1×10^−8^,1×10^−7^) combined with a baseline *μ* = 10^−8^. Starting at generation 10000 we sampled 200 individuals every 100 generations and calculated *F_ST_* between X and Y in 1kb windows. We ran three replicates of each scenario-parameter combination.

## Data accessibility

The genome assembly can be found at the NCBI SRA under BioProject accession: PRJNA1476984. Short read sequencing of *G. wheatlandi* samples collected in this study from the eight wild-caught *G. wheatlandi* individuals collected at Rainbow Haven Beach, Nova Scotia, Canada is currently being submitted to SRA and will be available via accession before final publication. Short read sequencing of *G. aculeatus* x *G. wheatlandi* crosses also used in this study can be found using accession: PRJNA742065. RNA seq data used for the genome annotation can be found at the accession PRJNA746773. The code for the empirical analyses in this study can be found at https://github.com/DanJeffries/Blackspotted_sex_chrom_recomb/ and code for the simulations can be found at https://github.com/wachalak/XY-divergence/.

## Acknowledgments

This work was supported by Swiss National Science Foundation grants 31003A_176130, 310030_204681, and TMAG-3_209309 to CLP. KW and SP were supported by the Swiss National Science Foundation project grant 10001034 awarded to SP.

## Author Contributions

DLJ, CLP and ZL conceived of the study. ZL and DLJ carried out all genomic analyses and interpretation with help from CLP and MK. KW and SP performed the simulations. MK and CC advised on the topology distribution analyses and interpretation. DLJ led the manuscript writing, with comments on manuscript drafts from all other authors.

## Supporting information

### Methods

#### Identification of centromeres

We first followed the approach of (100, see scripts published therein) to identify the most likely centromeric repeat monomer in the *G. wheatlandi* genome. We then aligned this sequence to our *G. wheatlandi* genome assembly to locate putative centromeric regions. Second, when performing the above-mentioned alignment, we noticed many alignments to unplaced scaffolds. We thus attempted to manually infer the location of these unplaced scaffolds by filtering for Hi-C read pairs in which one read aligns to an unplaced scaffold containing the centromeric repeat, and the other read aligns somewhere within a chromosome-level scaffold. Finally, we calculated population nucleotide diversity (*Pi*) in 10kb windows along the genome using the female samples in our WGS dataset. *Pi* is expected to decrease around centromeres owing to their lack of recombination, we used such regions of low Pi as additional evidence for the location of the centromere. As *Pi* is not informative for centromere positions in non-recombining regions this approach could not be used for the Y chromosome.

### Results

#### Pseudoautosomal regions on the sex chromosome assembly

We found an obvious PAR on Chr12 which we identified using male vs female coverage and *F*_ST_ analyses. We estimate this PAR is approximately 6 Mb in length. In contrast, we found no obvious evidence for a PAR on Chr19. The first genomic window containing SNPs on Chr19 is located around 64,000 bp from the end of the Chr19 X and has an F_ST_ of 0.115, which is already above the autosomal average. The proceeding windows along the chromosome have much higher F_ST_, in line with the rest of the sex-linked region of Chr19.

PARs on sex chromosomes are thought to be essential for ensuring normal segregation of chromosomes during meiosis. The Chr12 X and the Y can benefit from the PAR on Chr12 X, however, if indeed there is no PAR on the Chr19 X, it raises the question of how this chromosome properly segregates. In addition, (Sardell et al. 30) find a PAR on Chr19 in their analyses. Thus, we validated our result using several follow-up analyses. First, we aligned our Chr19 X assembly to the *G. aculeatus* assembly used in (originally from (Glazer et al. 101)) using NUCMER. We found no strong alignments in our *G. wheatlandi* Chr19 X assembly to the PAR in the *G. aculeatus*, including on unplaced scaffolds. Next, we aligned the raw PacBio sequencing used in our *G. wheatlandi* assembly to the *G. aculeatus* assembly. We found an extremely low depth of *G. wheatlandi* read alignments in the *G. aculeatus* PAR region. Together these two results suggest that the *G. wheatlandi* genome does not contain any region with good homology to the *G. aculeatus* PAR. We thus conclude that either there is no Chr19 PAR in *G. wheatlandi*, or that it is too small to be identified by our assembly.

Despite the above analyses, the question remains, why was a PAR on Chr19 X seen in the analyses in Sardell et al (30)? In that study, the authors produced *G. wheatlandi* x *G. aculeatus* F1 hybrids from *G. aculeatus* mothers and *G. wheatlandi* fathers. F1 hybrid female offspring thus had one Chr19 X from *G. aculeatus* and one from *G. wheatlandi*, while F1 hybrid male offspring had one Chr19 X from *G. aculeatus* and one *G. wheatlandi* Y chromosome. If a Chr19 X PAR exists in *G. aculeatus* but not in *G. wheatlandi*, then both F1 hybrid males and females would have one copy of this PAR on their *G. aculeatus* Chr19 X. When aligning hybrid sample reads to the *G. aculeatus* reference, reads from the PAR on the *G. aculeatus* Chr19 X in both sexes would align, while no reads from the *G. wheatlandi* Chr19 X or Y would align to this region. The ratio of read depth between males and females would thus be 1-1, despite the absence of this region on either the Chr19 X or Y in the *G. wheatlandi* genome.

#### Centromere positions

Centromeres were located using a combination of centromeric repeat sequence alignments to chromosome-level scaffolds, manual location of unplaced scaffolds with centromeric repeat alignments, and estimation of *Pi* along the genome (Figure S5). Alignments of centromeric repeats were found for 14 of the 21 chromosomes. In almost every case, these alignments were found in regions with depressed *Pi*, and high chance of containing an unplaced scaffold containing centromeric repeats, implying concordance between these approaches. On chromosomes 6 and 14, repeat alignments were found in two locations in the chromosome, a result that is biologically unlikely. However, in both cases, only one of the alignments was supported by *π* and the location of unplaced scaffolds containing centromeric sequences. We therefore took the location supported by all three approaches as our hypothesised centromere location in these cases. For the remaining 7 chromosomes for which no centromeric repeat alignment was found in the chromosome-level scaffolds, *π* and the likely placement of unplaced centromeric scaffolds were used to denote the centromere location.

While they do not identify all chromosomes in their study, our predictions of centromere locations agreed well with cytogenetic evidence from Ross et al. (27) in that the largest three chromosomes appear to be metacentric or submetacentric, chromosome 12 appears acrocentric or telocentric and the Y chromosome also presents as submetacentric.

Despite being the product of a fusion between two chromosomes, only one centromere could be found on the Y chromosome. This signal was located within the chromosome 19 region of the Y chromosome, however, it is most likely that this is the chromosome 12 centromere which has been successively repositioned by inversions that have occurred around the fusion point (see below).

#### LTR 2648

During repeat analyses of the sex chromosomes, we observed a surprising burst of insertions of a single repeat family, LTR_2648. Interestingly, our Kimura distance analyses of this family showed not only recent but also an old burst of insertions on the Y (around K2P = 20, see LTR panel for S6 in Figure S2). These results are somewhat confusing, as this region of the Y chromosome is relatively young, and K2P=20 likely predates its sex-linkage. However, could be explained by the erroneous categorisation of two cryptic LTR subfamilies within LTR_2648. The consistent divergence between copies from each subfamily would result in the bimodal peak observed. However, regardless of the presence of one or two types of LTR in this family, the Y specific burst of this family clearly contrasts with the lack of new repeat insertions from other families in S6.

The distribution of LTR_2648 repeats is also intriguing, being highly localised. We identified 257 copies of LTR_2648 on Chr12 X, concentrated in just two locations, ∼16Mb (S6) and ∼20Mb (S4). On chromosome Y the expansion of this repeat family has predominantly taken place at position ∼11.3Mb, which is homologous to the region at 16Mb within S6 on Chr12 X. The Y now hosts a total of 415 copies of LTR_2648, representing a 1.6 fold increase in copy number relative to the X. Importantly, this single repeat was also solely responsible for the young burst signal in this stratum.

#### Tandem repeats indicate fragile sites at the S6 breakpoints

Interestingly, the *F*_ST_ changepoints that delimit stratum S6 (Figure 2) align very closely to the breakpoints of the inversion that occurred in *G. wheatlandi* after divergence from *G. aculeatus* but before sex linkage evolved in the region (Figure S1). This prompted us to test for the frequency of tandem repeats in this region, which have been shown to associate with so-called “fragile sites” in the genome, *i.e.* sites that are prone to double strand breaks. We found that there is indeed increased tandem repeat density in the regions close to the S6 boundaries (Figure S2). 1kb windows at the left and right boundaries fall in the 82nd and 85th percentile of tandem repeat density per window respectively. However, this includes the tandem repeat rich PAR region. After removing this the windows left and right of S6 fall into the 92nd and 95th percentiles respectively. This result is consistent with the hypothesis that fragile sites could explain the shared breakpoints between these two structural rearrangement events.

### Discussion

#### On the biology of *G. wheatlandi* sex chromosomes

The rate of sex-linked inversion fixation and recombination loss on the Y in this species is seemingly rapid. We estimate that between 3-5 inversions have been fixed on the Chr12 region of the Y since the sex-chromosome autosome fusion occurred between 1.7M and 11.9M years ago (30). Further, based on synteny with closely related species, we find that the rearrangements between the X and Y observed here all occurred on the Y chromosome, despite theoretical predictions that X chromosomes should accumulate more inversions than Y chromosomes under models in which sexually antagonistic selection is the main driver of recombination suppression (102). This lack of inversions on the X thus raises the possibility that sexually antagonistic selection has not played an important role in the loss of recombination on *G. wheatlandi* sex chromosomes.

Another intriguing finding from our phased assemblies is that at least one inversion on the Y (Inversion D, Figure 1) has occurred within a preexisting inversion (Inversion C). Further, the extreme lack of synteny between the Chr19 X and Y region suggests that many rearrangements have occurred, some of which likely were likely nested within preexisting inversions. Why such nested inversions fix is an interesting question as presumably they have little effect on the recombination landscape, given that they occur in regions where recombination has already been suppressed. Thus, in such scenarios, models invoking beneficial linkage between the PSDL and nearby loci (e.g. to resolve sexual antagonism) do not apply. Might there be another benefit to sex-linked inversions on Y chromosomes other than their effect on recombination? One possibility could be that an inversion creates adaptive mutations at its breakpoints (103), for example by altering regulation of nearby genes, or perhaps by silencing genes which already have already accumulated deleterious mutations (17, 104). An alternative that should not be discounted however, is that such inversions are fixed simply via genetic drift and have no effect on fitness in either sex.

Aside from structural rearrangements, our phased assemblies also allowed us to identify the likely position of the centromere on the Y chromosome. Unfortunately, while π was very informative for the location of centromeres in autosomes, lack of recombination and sequence diversity on the Y precludes its use to infer the Y centromere. We were able to find a likely location for a Y centromere based on read pairs matching unplaced scaffolds containing centromere sequence, however we cannot distinguish whether this is the Chr12 or Chr19 centromere. The Chr19 centromere would likely have been weakened by mechanisms of Y degeneration by the time the fusion occurred, and thus it is most likely that the Chr12 centromere would be the one retained following the Y-autosome fusion (105). Further, the location of the identified centromere is consistent with the expected location of the Chr12 centromere based on the inferred sequence of rearrangements in this region. Still, we cannot definitively rule out the possibility that it is the Chr19 centromere.

One surprising result was the lack of an observable PAR on either the Chr19X or the Chr19 end of the Y chromosome. Chromosomes rely heavily on crossovers to aid proper segregation during meiosis (106). On sex chromosomes, such crossovers can only occur in the PAR, making these regions extremely important for proper sex chromosome segregation (107). The *G. wheatlandi* Y chromosome shares a PAR with Chr12X, thus these two chromosomes likely segregate properly. However, the lack of a PAR on Chr19X raises the question of how correct segregation is mediated on this chromosome. One possibility would be that the proper segregation of the Y, aided by the Chr12 PAR in turn facilitates the proper segregation of the Chr19X. However, another possibility would be that the Chr19 PAR is extremely small and was missed by our assembly and further analyses. In any case, it would also be interesting to test, across the multiple known XXY or ZZW systems created by sex chromosome to autosome fusions, how many retain PARs on both of the X or Z chromosomes?

A final intriguing observation from our genome assembly was the behaviour of the LTR_2648 repeat (Figure S2), which showed extremely high copy number, often in tandem, within the putatively recombined S6 stratum on Chr12 X and the homologous Y chromosome region. An obvious question leading from this result is whether this repeat family played a role in the recombination event that occurred in this region. While repeats have been found to be enriched at structural rearrangement break points (42), LTR_2648 is located well within the bounds of S6, making it unlikely that it contributed to the fragility of this region. However, the proliferation of repeats can be highly deleterious, and, while we cannot know whether this occurred prior to the X/Y recombination event at S6, the 1.6 fold increase in copies of this repeat on the present Y haplotype attests to its ability to proliferate once Y-linked. One hypothesis is therefore that selection favoured the recombinant Y haplotype due to the lower LTR_2648 copy number within S6. However, there is little possibility of testing this hypothesis.

### Supplementary figures

**Figure S1.**
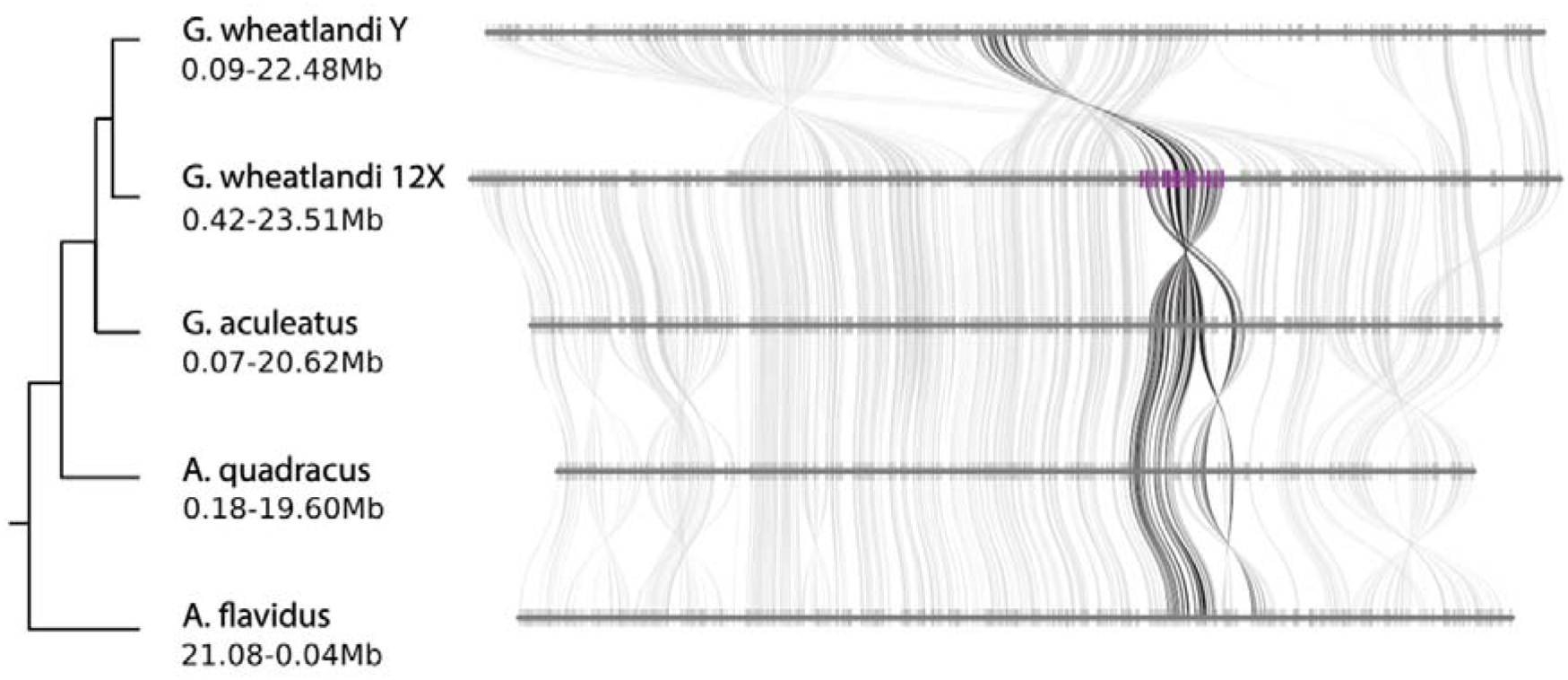
Chromosome 12 synteny of 1-to-1 orthologs between the *G. wheatlandi* and other stickleback species for which genome assemblies currently exist. Black gene links denote genes involved in a *G. wheatlandi* specific inversion on Chr12. Purple genes on the *G. wheatlandi* Chr12 X denote genes involved in the recombination event that created S6.

**Figure S2.**
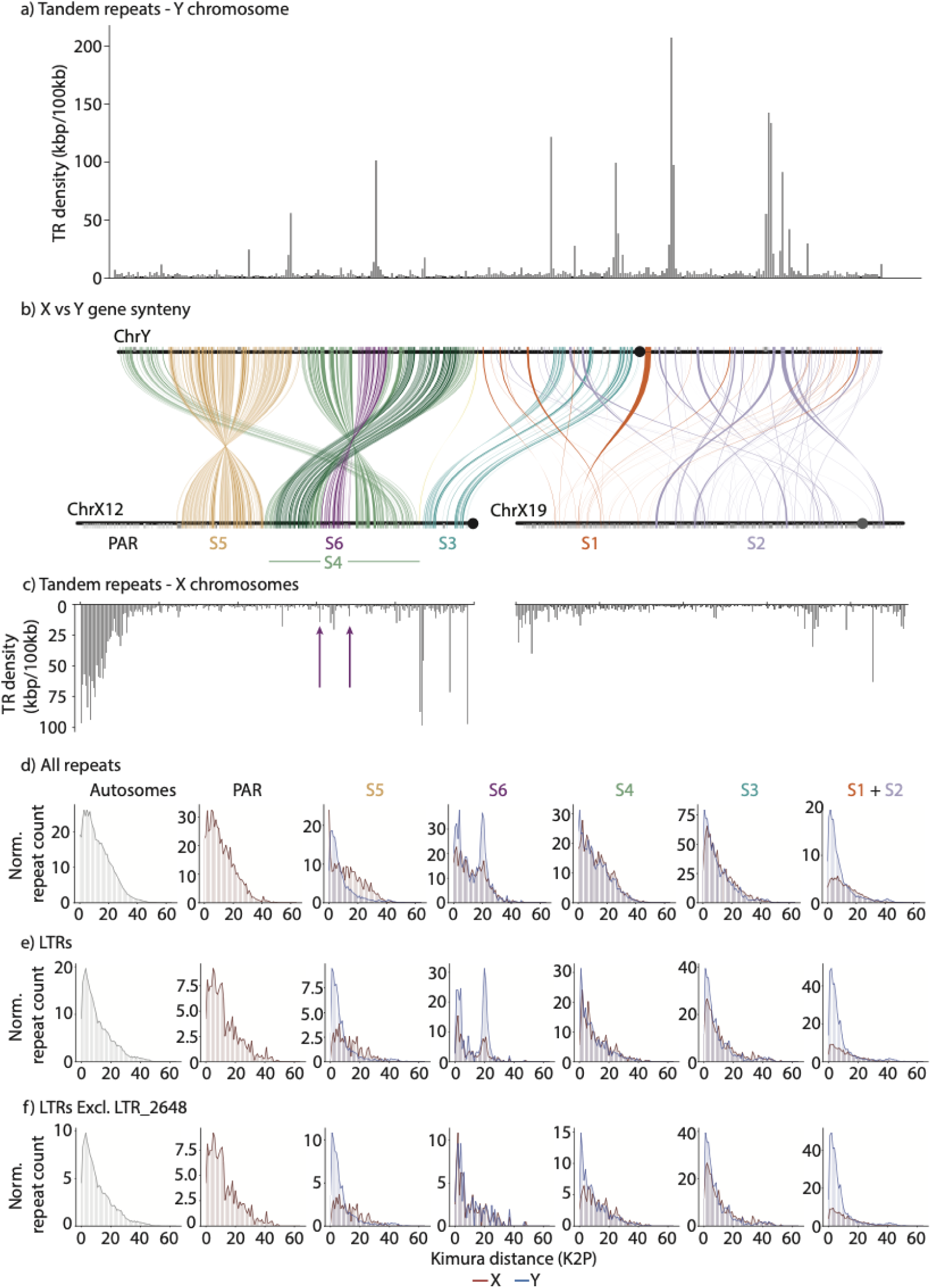
Repeat landscapes across the sex chromosomes. a) and c) show the density of tandem repeats on the Y and X chromosomes respectively. b) shows gene synteny differences between X and Y chromosomes as in Figure 2a. Remaining panels show repeat ages (Kimura distance (K2P)) for repeats of d) all families, e) LTRs only, and f) LTRs excluding LTR2648. Each plot shows the counts of repeats (Y axis) for a given K2P divergence (x axis) from the centroid sequence for its repeat family. Counts are normalised by the size of the stratum in which they are calculated to allow for comparisons across strata.

**Figure S3.**
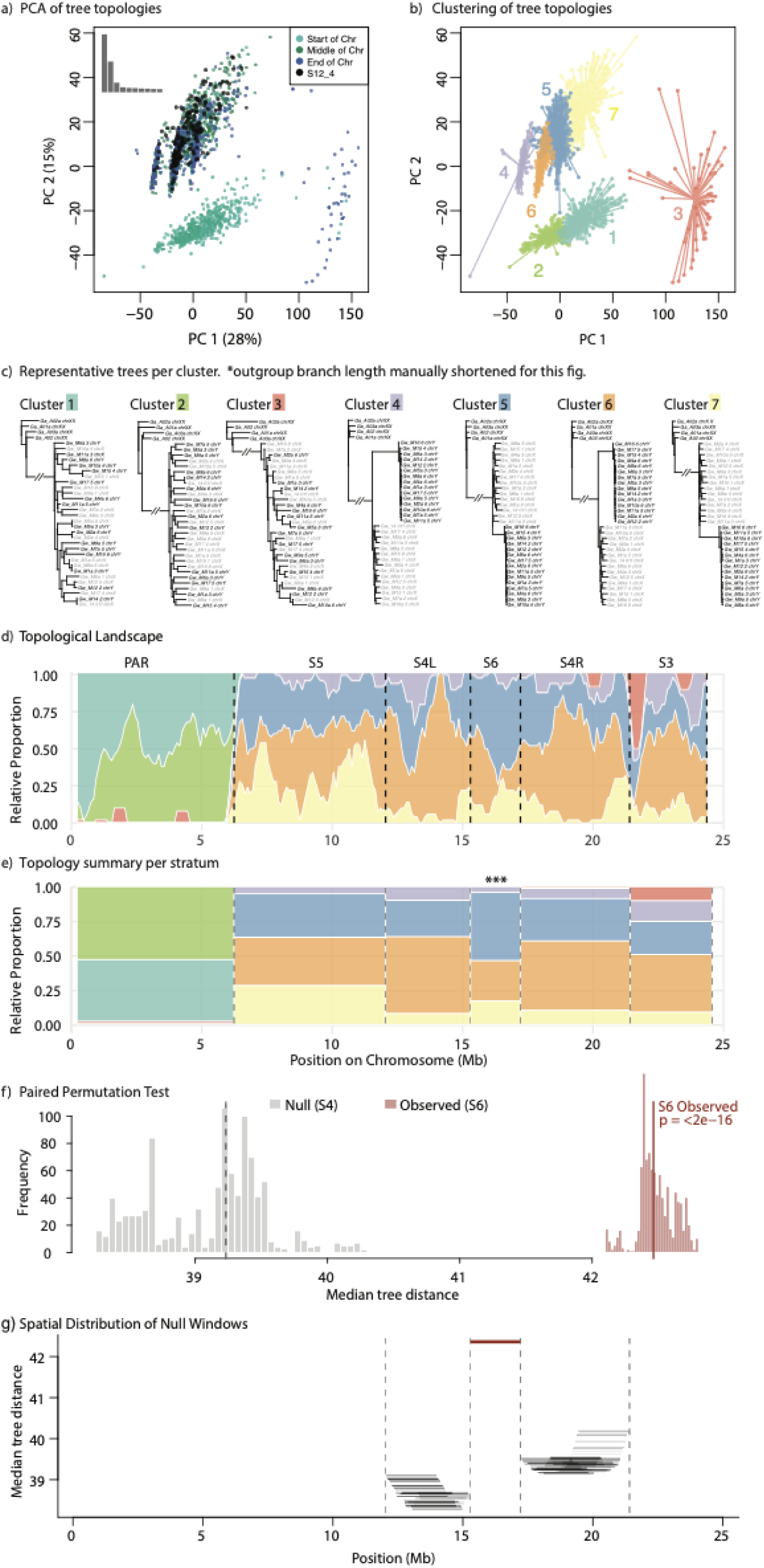
**a)** PCA of 2463 tree topologies, each from a 100kb window, with a 10kb step size (i.e. overlap) between windows. Points each represent a single tree and are coloured according to their position on Chr12. Black points represent topologies that fall within the putatively recombined stratum S6. **b)** Clusters of tree topology types based on the PCA in a). **c)** Representative trees (closest to cluster medians) for each topology cluster in b). G. aculeatus female (XX unphased) outgroup sequences denoted with “Ga”, phased *G. wheatlandi* Chr12X sequences (“*Gw*_<sampleID> chrX”) in grey, and phased *G. wheatlandi* Ch12Y sequences (“*Gw*_<sampleID> chrY”) in black. **d)** Relative proportions of tree topology clusters along Chr12, plotted in 5Mb windows (i.e. 50 trees per window) with 1Mb (10 tree) step size. **e)** Relative proportions of tree topology clusters summarised by stratum. Vertical dashed lines give stratum boundaries for reference. **f)** Null (grey) and observed (distribution) of tree distances calculated for topology permutation tests (see methods text for details). **g)** Spatial distribution of sampled topology windows, and their median distances to the reference background, used to create the null distribution in f). Location and median distance of S6 (test observation) shown in red.

**Figure S4.**
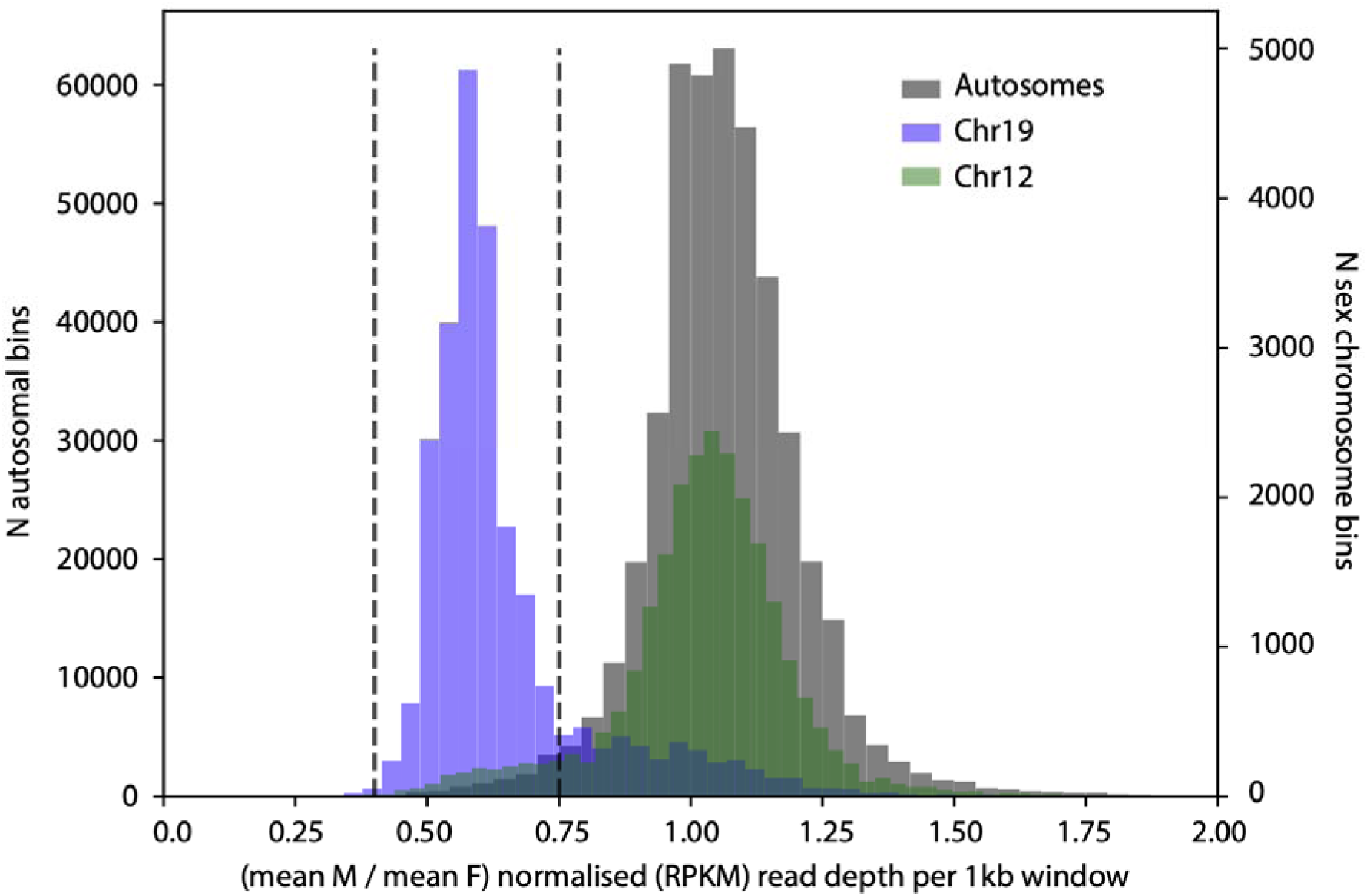
Histogram of mean male / mean female normalised (RPKM) read depth for 1kb windows, compared between autosomes (left Y axis) and the sex chromosomes (right Y axis). Dashed vertical lines show the thresholds used to assign windows as X-specific (see red points in Figure 2).

**Figure S5.**
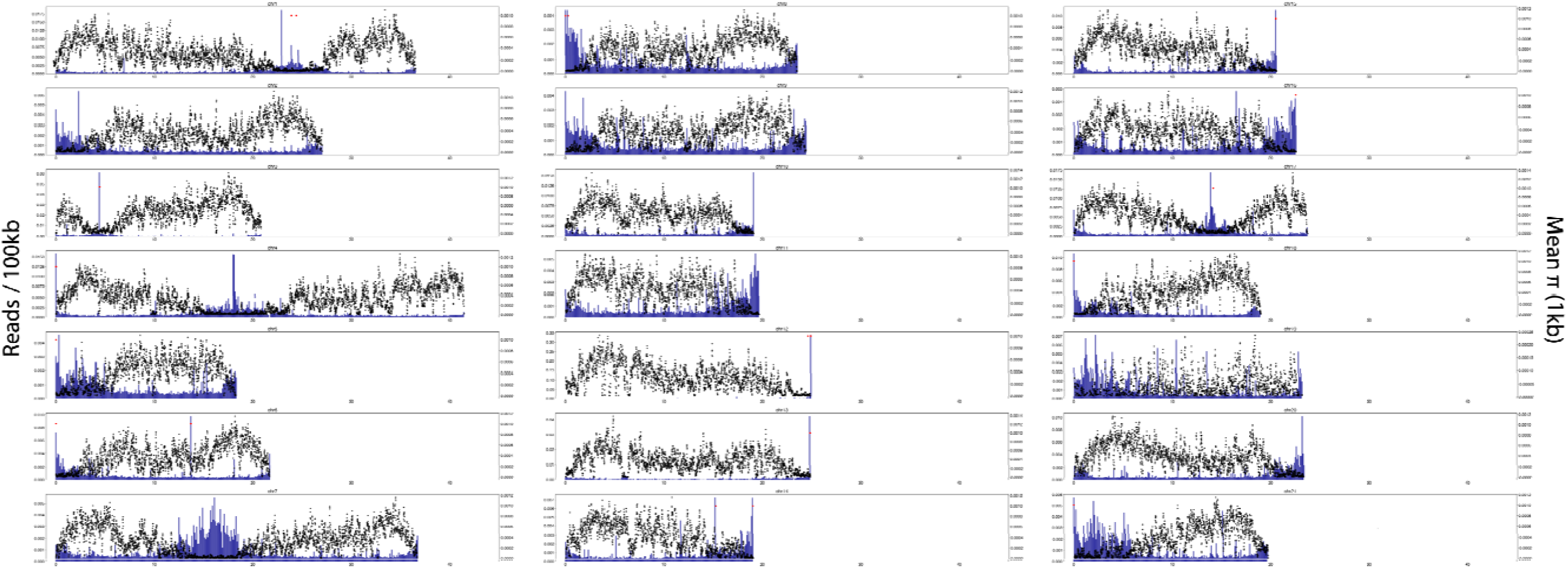
Location of centromeric regions. Blue bars (left Y axis) represent the proportion of reads in each 100kb window in a chromosome with mates hitting an unplaced centromere scaffold. Black points (right Y axis) represent average π calculated in 1kb bins from female WGS data. Red points (arbitrary Y coordinates) denote positions of centromeric repeat sequence alignments.

## Appendix 1

GIF animation of the hypothesised sequence of inversions and the recombination event leading to the observed contemporary gene order and differentiation on the neo-Y chromosome. For initial submission, download file here and view in any internet browser: https://www.dropbox.com/scl/fi/utgdzgfvgujoxluj2rcmv/Appendix_1_secondary_recomb_animation.gif?rlkey=n8tlqzoszqyj1sqxunh7zohtz&st=6wkm8mi4&dl=0

## Appendix 2

GIF animation of the forward simulations performed in SLiM of an evolving sex chromosome with a 5Mb non-recombining region denoted by “Breakpoints” and with gene flux possible within this region via double crossovers between X and Y. The simulated non-recombining region contained either a PSDL only (left panels), or PSDL + Sexually Antagonistic Loci (SAL) (right panels), and simulations were conducted for three combinations of baseline mutation and recombination rates (rows). X vs Y *F*_ST_ was calculated in 1kb windows, and the rolling Mean *F*_ST_ was calculated using 10 windows with a 1 window step. Each time step shows X vs Y *F*_ST_ every 100 generations. For initial submission, download file here and view in any internet browser: https://www.dropbox.com/scl/fi/pieki3mdftqhy94atbiuq/Appendix_2_XYevosims.gif?rlkey=0fu1reousoq0z3mivw701lafv&st=4×777yf3&dl=0

## References

1. D. Charlesworth, B. Charlesworth, G. Marais, Steps in the evolution of heteromorphic sex chromosomes. Heredity 95, 118–128 (2005).

2. B. A. Sandkam, et al., Extreme Y chromosome polymorphism corresponds to five male reproductive morphs of a freshwater fish. Nature Ecology & Evolution 5, 939–948 (2021).

3. A. P. Arnold, Y chromosome’s roles in sex differences in disease. Proc. Natl. Acad. Sci. U. S. A. 114, 3787–3789 (2017).

4. T. Connallon, et al., Local adaptation and the evolution of inversions on sex chromosomes and autosomes. Philos. Trans. R. Soc. Lond. B Biol. Sci. 373, 20170423 (2018).

5. J. Kitano, et al., A role for a neo-sex chromosome in stickleback speciation. Nature 461, 1079–1083 (2009).

6. J. B. S. Haldane, Sex ratio and unisexual sterility in hybrid animals. J. Genet. 12, 101–109 (1922).

7. L. F. Delph, J. P. Demuth, Haldane’s rule: genetic bases and their empirical support. J. Hered. 107, 383–391 (2016).

8. D. C. Presgraves, Evaluating genomic signatures of “the large X-effect” during complex speciation. Mol. Ecol. 27, 3822–3830 (2018).

9. P. Jay, D. Jeffries, F. E. Hartmann, A. Véber, T. Giraud, Why do sex chromosomes progressively lose recombination? Trends Genet. 40, 564–579 (2024).

10. S. Ponnikas, H. Sigeman, J. K. Abbott, B. Hansson, Why Do Sex Chromosomes Stop Recombining? Trends Genet. 34, 492–503 (2018).

11. D. Cortez, R. Marin, D. Toledo-Flores, L. Froidevaux, Origins and functional evolution of Y chromosomes across mammals. Nature (2014).

12. E. D. Jarvis, et al., Whole-genome analyses resolve early branches in the tree of life of modern birds. Science 346, 1320–1331 (2014).

13. Q. Zhou, et al., Complex evolutionary trajectories of sex chromosomes across bird taxa. Science 346, 1246338 (2014).

14. K. Sahara, A. Yoshido, W. Traut, Sex chromosome evolution in moths and butterflies. Chromosome Res. 20, 83–94 (2012).

15. D. Bachtrog, The temporal dynamics of processes underlying Y chromosome degeneration. Genetics 179, 1513–1525 (2008).

16. T. Lenormand, D. Roze, Can mechanistic constraints on recombination reestablishment explain the long-term maintenance of degenerate sex chromosomes? Peer Community J. 4 (2024).

17. T. Lenormand, D. Roze, Y recombination arrest and degeneration in the absence of sexual dimorphism. Science 375, 663–666 (2022).

18. C. Grossen, S. Neuenschwander, N. Perrin, The evolution of XY recombination: sexually antagonistic selection versus deleterious mutation load. Evolution 66, 3155–3166 (2012).

19. O. Blaser, S. Neuenschwander, N. Perrin, Sex-chromosome turnovers: the hot-potato model. Am. Nat. 183, 140–146 (2014).

20. O. Blaser, C. Grossen, S. Neuenschwander, N. Perrin, Sex-chromosome turnovers induced by deleterious mutation load. Evolution 67, 635–645 (2013).

21. C. L. Peichel, et al., Assembly of the threespine stickleback Y chromosome reveals convergent signatures of sex chromosome evolution. Genome Biol. 21, 177 (2020).

22. C. Lemaitre, et al., Footprints of inversions at present and past pseudoautosomal boundaries in human sex chromosomes. Genome Biol. Evol. 1, 56–66 (2009).

23. C. Moraga, et al., The *Silene latifolia* genome and its giant Y chromosome. Science 387, 630–636 (2025).

24. H. P. Yazdi, H. Ellegren, A genetic map of ostrich Z chromosome and the role of inversions in avian sex chromosome evolution. Genome Biol. Evol. 10, 2049–2060 (2018).

25. P. Jay, E. Tezenas, A. Véber, T. Giraud, Sheltering of deleterious mutations explains the stepwise extension of recombination suppression on sex chromosomes and other supergenes. PLoS Biol. 20, e3001698 (2022).

26. X. Cheng, et al., Evolution of a ZW sex determination system in sticklebacks. Sci. Adv. 12, eaeb1484 (2026).

27. J. A. Ross, J. R. Urton, J. Boland, M. D. Shapiro, C. L. Peichel, Turnover of sex chromosomes in the stickleback fishes (gasterosteidae). PLoS Genet. 5, e1000391 (2009).

28. Z. Liu, et al., The fourspine stickleback (Apeltes quadracus) has an XY sex chromosome system with polymorphic inversions on both X and Y chromosomes. PLoS Genet. 21, e1011465 (2025).

29. D. L. Jeffries, J. A. Mee, C. L. Peichel, Identification of a candidate sex determination gene in *Culaea inconstans* suggests convergent recruitment of an Amh duplicate in two lineages of stickleback. J. Evol. Biol. 35, 1683–1695 (2022).

30. J. M. Sardell, M. P. Josephson, A. C. Dalziel, C. L. Peichel, M. Kirkpatrick, Heterogeneous histories of recombination suppression on stickleback sex chromosomes. Mol. Biol. Evol. 38, 4403–4418 (2021).

31. S. Varadharajan, et al., A high-quality assembly of the nine-spined stickleback (*Pungitius pungitius*) genome. Genome Biol. Evol. 11, 3291–3308 (2019).

32. T. R. Chen, H. M. Reisman, A comparative chromosome study of the North American species of sticklebacks (Teleostei: Gasterosteidae). Cytogenetics 9, 321–332 (1970).

33. R. Hobza, et al., Impact of repetitive DNA on sex chromosome evolution in plants. Chromosome Res. 23, 561–570 (2015).

34. V. Peona, et al., The avian W chromosome is a refugium for endogenous retroviruses with likely effects on female-biased mutational load and genetic incompatibilities. Philos. Trans. R. Soc. Lond. B Biol. Sci. 376, 20200186 (2021).

35. M. B. Cioffi, J. P. M. Camacho, L. A. C. Bertollo, Repetitive DNAs and differentiation of sex chromosomes in neotropical fishes. Cytogenet. Genome Res. 132, 188–194 (2011).

36. L. S. Stevison, K. B. Hoehn, M. A. F. Noor, Effects of inversions on within- and between-species recombination and divergence. Genome Biol. Evol. 3, 830–841 (2011).

37. K. N. Crown, D. E. Miller, J. Sekelsky, R. S. Hawley, Local inversion heterozygosity alters recombination throughout the genome. Current Biology 28, 2984–2990 (2018).

38. R. De-Kayne, et al., Incomplete recombination suppression fuels extensive haplotype diversity in a butterfly colour pattern supergene. PLoS Biol. 23, e3003043 (2025).

39. M. Matschiner, et al., Supergene origin and maintenance in Atlantic cod. *Nat*. Ecol. Evol. 6, 469–481 (2022).

40. M. Rönspies, et al., Massive crossover suppression by CRISPR-Cas-mediated plant chromosome engineering. Nat. Plants 8, 1153–1159 (2022).

41. T. Decroly, R. Vila, K. Lohse, A. Mackintosh, Rewinding the Ratchet: Rare Recombination Locally Rescues Neo-W Degeneration and Generates Plateaus of Sex-Chromosome Divergence. Mol. Biol. Evol. 41 (2024).

42. A. Ruiz-Herrera, J. Castresana, T. J. Robinson, Is mammalian chromosomal evolution driven by regions of genome fragility? Genome Biol. 7, R115 (2006).

43. D. Charlesworth, The timing of genetic degeneration of sex chromosomes. Philos. Trans. R. Soc. Lond. B Biol. Sci. 376, 20200093 (2021).

44. C. Schmidt, et al., Changing local recombination patterns in Arabidopsis by CRISPR/Cas mediated chromosome engineering. Nat. Commun. 11, 1–8 (2020).

45. A. Navarro, E. Betrán, A. Barbadilla, A. Ruiz, Recombination and gene flux caused by gene conversion and crossing over in inversion heterokaryotypes. Genetics 146, 695–709 (1997).

46. R. F. Guerrero, F. Rousset, M. Kirkpatrick, Coalescent patterns for chromosomal inversions in divergent populations. Philos. Trans. R. Soc. Lond. B Biol. Sci. 367, 430–438 (2012).

47. C. Cheng, et al., Ecological genomics of *Anopheles gambiae* along a latitudinal cline: a population-resequencing approach. Genetics 190, 1417–1432 (2012).

48. M. Kapun, E. D. Mitchell, T. J. Kawecki, P. Schmidt, T. Flatt, An ancestral balanced inversion polymorphism confers global adaptation. Mol. Biol. Evol. 40, msad118 (2023).

49. F. Ruzicka, et al., The search for sexually antagonistic genes: Practical insights from studies of local adaptation and statistical genomics. Evol Lett 4, 398–415 (2020).

50. A. J. Dagilis, et al., Searching for signatures of sexually antagonistic selection on stickleback sex chromosomes. Philos. Trans. R. Soc. Lond. B Biol. Sci. 377, 20210205 (2022).

51. M. Kirkpatrick, R. F. Guerrero, S. V. Scarpino, Patterns of neutral genetic variation on recombining sex chromosomes. Genetics 184, 1141–1152 (2010).

52. K. A. Behrens, et al., A chromosome inversion creates a supergene for sex and colour in Lake Malawi cichlids. Mol. Ecol. 34, e17821 (2025).

53. J. N. Brand, et al., Stepwise emergence of recombination suppression precedes fissiparous asexuality in the planarian *Schmidtea mediterranea*. Nat. Commun. 17, 5588 (2026).

54. C. Zhang, et al., De Novo Mutation Rates in Sticklebacks. Mol. Biol. Evol. 40 (2023).

55. C. Zhang, et al., Rate of de novo mutations in the three-spined stickleback. Heredity 134, 387–395 (2025).

56. T. Lenormand, D. Roze, A single theory for the evolution of sex chromosomes and the two rules of speciation. Science 389, eado9032 (2025).

57. Z. Liu, et al., Chromosomal fusions facilitate adaptation to divergent environments in threespine stickleback. Mol. Biol. Evol. 39 (2022).

58. X.-B. Wang, et al., An effective strategy for assembling the sex-limited chromosome. Gigascience 13 (2024).

59. H. Li, Minimap2: pairwise alignment for nucleotide sequences. Bioinformatics 34, 3094–3100 (2018).

60. F. Ramírez, et al., deepTools2: a next generation web server for deep-sequencing data analysis. Nucleic Acids Res. 44, W160–5 (2016).

61. M. Kolmogorov, J. Yuan, Y. Lin, P. A. Pevzner, Assembly of long, error-prone reads using repeat graphs. Nat. Biotechnol. 37, 540–546 (2019).

62. R. Vaser, I. Sović, N. Nagarajan, M. Šikić, Fast and accurate de novo genome assembly from long uncorrected reads. Genome Res. 27, 737–746 (2017).

63. B. J. Walker, et al., Pilon: an integrated tool for comprehensive microbial variant detection and genome assembly improvement. PLoS One 9, e112963 (2014).

64. D. Guan, et al., Identifying and removing haplotypic duplication in primary genome assemblies. Bioinformatics 36, 2896–2898 (2020).

65. H. Zhang, et al., Fast alignment and preprocessing of chromatin profiles with Chromap. Nat. Commun. 12, 6566 (2021).

66. C. Zhou, S. A. McCarthy, R. Durbin, YaHS: yet another Hi-C scaffolding tool. Bioinformatics 39 (2023).

67. M. Xu, et al., TGS-GapCloser: A fast and accurate gap closer for large genomes with low coverage of error-prone long reads, GigaScience. 9 (2020).

68. M. Manni, M. R. Berkeley, M. Seppey, F. A. Simão, E. M. Zdobnov, BUSCO update: Novel and streamlined workflows along with broader and deeper phylogenetic coverage for scoring of eukaryotic, prokaryotic, and viral genomes. Mol. Biol. Evol. 38, 4647–4654 (2021).

69. S. Ou, et al., Benchmarking transposable element annotation methods for creation of a streamlined, comprehensive pipeline. Genome Biol. 20, 275 (2019).

70. H. Yan, A. Bombarely, S. Li, DeepTE: a computational method for de novo classification of transposons with convolutional neural network. Bioinformatics 36, 4269–4275 (2020).

71. A. F. A. Smit, R. Hubley, P. Green, RepeatMasker (2013).

72. A. Dobin, et al., STAR: ultrafast universal RNA-seq aligner. Bioinformatics 29, 15–21 (2013).

73. L. Gabriel, et al., BRAKER3: Fully automated genome annotation using RNA-seq and protein evidence with GeneMark-ETP, AUGUSTUS, and TSEBRA. Genome Res. 34, 769–777 (2024).

74. J. Huerta-Cepas, et al., Fast genome-wide functional annotation through orthology assignment by eggNOG-mapper. Mol. Biol. Evol. 34, 2115–2122 (2017).

75. G. Marçais, et al., MUMmer4: A fast and versatile genome alignment system. PLoS Comput. Biol. 14, e1005944 (2018).

76. J. Dainat, AGAT: Another Gff Analysis Toolkit to handle annotations in any GTF/GFF format (2022).

77. H. Tang, et al., Synteny and collinearity in plant genomes. Science 320, 486–488 (2008).

78. W. H. Gates, C. H. Papadimitriou, Bounds for sorting by prefix reversal. Discrete Math. 27, 47–57 (1979).

79. G. Tesler, GRIMM: genome rearrangements web server. Bioinformatics 18, 492–493 (2002).

80. D. Wang, L. Wang, GRSR: a tool for deriving genome rearrangement scenarios from multiple unichromosomal genome sequences. BMC Bioinformatics 19, 291 (2018).

81. S. Andrews, FastQC: a quality control tool for high throughput sequence data (Babraham Bioinformatics, 2010).

82. A. M. Bolger, M. Lohse, B. Usadel, Trimmomatic: a flexible trimmer for Illumina sequence data. Bioinformatics 30, 2114–2120 (2014).

83. H. Li, R. Durbin, Fast and accurate short read alignment with Burrows-Wheeler transform. Bioinformatics 25, 1754–1760 (2009).

84. H. Li, et al., The Sequence Alignment/Map format and SAMtools. Bioinformatics 25, 2078–2079 (2009).

85. A. Tarasov, A. J. Vilella, E. Cuppen, I. J. Nijman, P. Prins, Sambamba: fast processing of NGS alignment formats. Bioinformatics 31, 2032–2034 (2015).

86. G. A. Van der Auwera, B. D. O’Connor, Genomics in the Cloud: Using Docker, GATK, and WDL in Terra (“O’Reilly Media, Inc.,” 2020).

87. P. Danecek, et al., The variant call format and VCFtools. Bioinformatics 27, 2156–2158 (2011).

88. C. Truong, L. Oudre, N. Vayatis, Selective review of offline change point detection methods. Signal Processing 167, 107299 (2020).

89. D. M. Emms, S. Kelly, OrthoFinder: phylogenetic orthology inference for comparative genomics. Genome Biol. 20, 238 (2019).

90. D. Wang, et al., Improved assembly of the *Pungitius pungitius* reference genome. G3 (Bethesda) 14 (2024).

91. A. Löytynoja, N. Goldman, An algorithm for progressive multiple alignment of sequences with insertions. Proc. Natl. Acad. Sci. U. S. A. 102, 10557–10562 (2005).

92. M. Suyama, D. Torrents, P. Bork, PAL2NAL: robust conversion of protein sequence alignments into the corresponding codon alignments. Nucleic Acids Res. 34, W609–12 (2006).

93. Z. Zhang, KaKs_Calculator 3.0: Calculating selective pressure on coding and non-coding sequences. Genomics Proteomics Bioinformatics 20, 536–540 (2022).

94. P. Danecek, et al., Twelve years of SAMtools and BCFtools. Gigascience 10 (2021).

95. T. Jombart, M. Kendall, J. Almagro-Garcia, C. Colijn, treespace: Statistical exploration of landscapes of phylogenetic trees. Mol. Ecol. Resour. 17, 1385–1392 (2017).

96. M. Kendall, C. Colijn, Mapping phylogenetic trees to reveal distinct patterns of evolution. Mol. Biol. Evol. 33, 2735–2743 (2016).

97. S. Seabold, J. Perktold, Statsmodels: Econometric and statistical modeling with python in Proceedings of the Python in Science Conference, (SciPy, 2010), pp. 92–96.

98. B. C. Haller, P. L. Ralph, P. W. Messer, SLiM 5: Eco-evolutionary simulations across multiple chromosomes and full genomes. Mol. Biol. Evol. 43, msaf313 (2026).

99. J. H. Gillespie, Population Genetics: A Concise Guide, 2nd Ed. (Johns Hopkins University Press, 2004).

100. D. P. Melters, et al., Comparative analysis of tandem repeats from hundreds of species reveals unique insights into centromere evolution. Genome Biol. 14, R10 (2013).

101. A. M. Glazer, E. E. Killingbeck, T. Mitros, D. S. Rokhsar, C. T. Miller, Genome assembly improvement and mapping convergently evolved skeletal traits in sticklebacks with genotyping-by-sequencing. G3 (Bethesda) 5, 1463–1472 (2015).

102. E. Flintham, C. Mullon, Sexual antagonism, mating systems, and recombination suppression on sex chromosomes. Evol. Lett. qrag021 (2026).

103. R. Villoutreix, et al., Inversion breakpoints and the evolution of supergenes. Mol. Ecol. 30, 2738–2755 (2021).

104. T. Lenormand, F. Fyon, E. Sun, D. Roze, Sex Chromosome Degeneration by Regulatory Evolution. Curr Biol (2020). 10.1016/j.cub.2020.05.052.

105. J. N. Cech, C. L. Peichel, Centromere inactivation on a neo-Y fusion chromosome in threespine stickleback fish. Chromosome Res. 24, 437–450 (2016).

106. Y. Hirose, et al., Chiasmata promote monopolar attachment of sister chromatids and their co-segregation toward the proper pole during meiosis I. PLoS Genet. 7, e1001329 (2011).

107. L. Kauppi, et al., Distinct properties of the XY pseudoautosomal region crucial for male meiosis. Science 331, 916–920 (2011).

